# Transcription-coupled structural dynamics of topologically associating domains regulate replication origin efficiency

**DOI:** 10.1101/2020.08.16.252668

**Authors:** Yongzheng Li, Boxin Xue, Liwei Zhang, Qian Peter Su, Mengling Zhang, Haizhen Long, Yao Wang, Yanyan Jin, Yingping Hou, Yuan Cao, Guohong Li, Yujie Sun

## Abstract

Metazoan cells only utilize a small subset of the potential DNA replication origins to duplicate the whole genome in each cell cycle. Origin choice is linked to cell growth, differentiation, and replication stress. Despite various genetic and epigenetic signatures are found to be related with active origins, it remains elusive how the selection of origins is determined. The classic Rosette model proposes that the origins clustered in a chromatin domain are preferentially and simultaneously fired, but direct imaging evidence has been lacking due to insufficient spatial resolution. Here, we applied dual-color stochastic optical reconstruction microscopy (STORM) super-resolution imaging to map the spatial distribution of origins within individual topologically associating domains (TADs). We found that multiple replication origins initiate separately at the spatial boundary of a TAD at the beginning of the S phase, in contrary to the Rosette model. Intriguingly, while both active and dormant origins are distributed homogeneously in the TAD during the G1 phase, active origins relocate to the TAD periphery before entering the S phase. We proved that such origin relocalization is dependent on both transcription and CTCF-mediated chromatin structure. Further, we observed that the replication machinery protein PCNA forms immobile clusters around the TADs at the G1/S transition, which explains why origins at the TAD periphery are preferentially fired. Thus, we propose a “Chromatin Re-organization Induced Selective Initiation” (CRISI) model that the transcription-coupled chromatin structural re-organization determines the selection of replication origins, which transcends the scope of specific genetic and epigenetic signatures for origin efficiency. Our *in situ* super-resolution imaging unveiled coordination among DNA replication, transcription, and chromatin organization inside individual TADs, providing new insights into the biological functions of sub-domain chromatin structural dynamics.

## INTRODUCTION

DNA replication is an exquisitely regulated process. Its deregulation may lead to genome instability and tumorigenesis [1]. In metazoans, duplication of the genome is initiated at tens of thousands of discrete chromosome loci known as replication origins. Intriguingly, while a mammalian cell has a total of ∼250,000 potential replication origins, it only uses a small subset (∼10%) to duplicate the whole genome [2–5]. Dormant origins can be initiated to rescue stalled replication forks, facilitating accurate and timely genome replication under stress [6, 7].

Whether the selection of origins is random or regulated has been under debate. Single cell-based measurements, including the classic DNA combing assays [8–10] and the recent single-cell sequencing studies [11, 12], showed that cells rarely use the same cohort of origins to duplicate the genome. Nevertheless, either single cell or population-averaged origin mapping experiments have confirmed that not all origins are equal, and they rather have differential probabilities of firing, namely origin efficiency [4], against a random origin selection mechanism.

The mechanisms that determine the origin efficiency remain enigmatic. Various genetic and epigenetic signatures, including CpG islands, G-quadruplexes, nucleosome-depleted regions, and histone modifications, are found to correlate with origin efficiency, but a consensus principle is still lacking [2]. The origin efficiency has been also suggested to link with chromatin structures [4, 13, 14]. Several earlier studies have revealed a relationship between replicons and chromatin loops [9, 15–17]. Beyond the loop structure, more recent studies have shown that the spatiotemporal initiation of replication is regulated at the chromatin domain level. Genome-scale mapping of replication kinetics showed that DNA replication in metazoan cells takes place in a defined temporal order with the genome segmented into large chromosomal regions, known as replication domains (RDs) [4, 5]. Each RD contains multiple replicons with uniform and constant replication timing. Importantly, the boundaries of RDs are found to align precisely with that of topologically associating domains (TADs) [18]. TADs are physical compartmentalization units of the genome that are stable over multiple cell cycles and conserved across related species [19]. Thus, this finding provides strong supports for the correlation between DNA replication and the three-dimensional (3D) structure of chromosomes. A typical RD is about 1 Mb and contains a few dozens of potential origins. These origins do not have similar replication efficiencies as only several of them are actually used to replicate the domain [2–5]. In temporal space, direct measurements on spread-out DNA fibers by DNA combing experiments have shown that the active origins within a RD fire nearly synchronously [8–10]. However, how the origin efficiency is spatially regulated in a RD has been an outstanding question.

In physical space, mapping of the spatial arrangements of replication sites by *in situ* fluorescence imaging in the nucleus showed that DNA replication initiates at thousands discrete puncta termed replication foci (RFi) [9, 20–25]. Provided that the number of pulse-labelled RFi is much less than the number of active origins [9, 20–22], RFi are thought to be the equivalents of RDs defined in temporal space and contain multiple replicons. Based on the collective evidence from the DNA halo [9, 15–17], DNA combing [8, 9, 26] and RFi imaging [9, 20–22], the Rosette model was proposed to illustrate the spatiotemporal organization and regulation of active origins in a RD [4]. In this model, a RD contains multiple loops which form a rosette-like structure with the active origins clustered and co-fired in the chromatin domain. The Rosette model is further supported by another study showing that depletion of cohesin, a protein complex scaffolding the rosette-like structure, reduces the number of origins used for genome duplication [27]. However, as previous imaging studies were mostly limited in their spatial resolution and lack of sequence-specific labeling of TADs and replication origins, there is no direct imaging evidence to support the clustered origin distribution. In a recent work, Cardoso and his colleagues applied SIM super-resolution imaging (resolution ∼100 nm) and identified more replicons than conventional imaging (resolution ∼300 nm) [23]. Importantly, with the improvement in resolution, nearly all replicons are found to be spatially separated at the beginning of the S phase, which casts doubt on the clustering of replication origins proposed in the Rosette model. In the accompanying work, they proposed a stochastic, proximity-induced (domino-like) replication initiation model, in which the active origins are not necessarily clustered spatially in the domain but the domino-like replication progression leads to clustering of replicons [28].

A thorough understanding of how the physical structure of RDs regulates origin efficiency needs *in situ* imaging of the spatial distribution of both active and dormant origins within the TADs. Given that a TAD is about 800 kb [18] with a radius of gyration less than 300 nm [29] and contains a few dozens of potential origins, dual-color 3D super-resolution imaging with ultra-high resolution in all three dimensions is a pre-requisite to distinguish which origins are more preferentially used among the many potential candidates within individual TADs. Moreover, new labelling strategies are necessary for dormant origins because the traditional approach based on metabolic pulse-labeling only labels active origins. Here, we applied a recently developed chromatin painting and imaging technique, namely OligoSTORM [30, 31], to investigate whether origin efficiency is dependent on TAD structure. OligoSTORM combines Oligopaints [32] with stochastic optical reconstruction microscopy (STORM) [33]. Oligopaints are highly efficient oligonucleotide FISH (fluorescent in situ hybridization) probes based on PCR strategy that can robustly label whole chromosomes or any specific chromosomal regions. STORM and its equivalents PALM/fPALM [34, 35] are single-molecule localization-based super-resolution imaging techniques that have the highest spatial resolution (∼20 nm laterally and ∼50 nm axially) among all super-resolution imaging methods [36]. With the best of both sides, OligoSTORM has been successfully applied to resolve the fine physical chromatin structures, such as TADs and compartments, in single cells [29, 37, 38].

Using OligoSTORM, we performed, to our knowledge, the first quantitative characterization of TAD structural dynamics and the spatiotemporal distribution of replication origins within individual TADs in different cell cycle stages at sub-diffraction-limit resolution. We discovered that in the G1 phase, TADs undergo a transcription-dependent structural re-organization process, which exposes active origins to the spatial boundary of the TAD; in contrast, dormant origins remain inside the TAD. We also observed an interesting peri-RFi distribution of the major replication machinery protein PCNA, in line with the observation that replication initiation generally takes place at the spatial boundary of a TAD. Thus, our work reveals a new origin selection mechanism that the replication efficiency of origins is determined by their physical distribution in the chromatin domain and transcription plays a role in the chromatin structural re-organization. This new mechanism transcends the scope of specific genetic and epigenetic signatures for origin efficiency and also provides new insight into the biological functions of sub-domain chromatin structural dynamics.

## RESULTS

### Replication origins initiate separately at the periphery of a TAD

In order to investigate the role of chromatin structure in origin selection, we chose to directly visualize how replication initiation is spatially organized and regulated within individual RDs using STORM imaging. Two RDs were chosen from the replication timing profile of HeLa cells (Appendix Figure S1a). The boundaries of either RD are overlaid with that of a TAD, which are hereafter named as TAD1 and TAD2, respectively (Appendix Figure S1b). TAD1 (Chr1:16911932-17714928) is an early replicating domain and enriched of transcriptionally active histone modifications. TAD2 (Chr1: 17722716-18846245) is a middle replicating domain and enriched of transcriptionally repressed histone modifications (Appendix Figure S1c). The two TADs were labeled by the Oligopaint approach using 12,000 TAMRA-modified primary oligonucleotide probes targeting the TADs and then imaged by STORM (Methods). Morphological characterization showed that the radii of gyration of TAD1 and TAD2 are about 200 nm (Appendix Figure S2), which is consistent with previous work [29, 38]. Moreover, even though the genomic length of TAD2 is larger than that of TAD1, the physical size of TAD2 is significantly smaller than that of TAD1 (Appendix Figure S2b), suggesting that TAD2 is more compact. This observation is consistent with the previously reported findings that active chromatin domains are more open than repressed chromatin domains [29, 38], thereby benchmarking the technical rigor of our TAD labeling and imaging.

Next, to visualize the replication initiation sites in the TADs, we took the classic metabolic pulse-labeling strategy [9]. Briefly, we first synchronized HeLa cells to the boundary of the G1 phase and S phase as previously described [9, 39] (Methods). Immediately after release of aphidicolin arrest at the G1/S boundary, we performed a 10-min pulse labeling of the replicating DNA by supplying thymidine analog EdU, which was then labeled with Alexa647 by click chemistry after fixation of the cell (Methods). This synchronization procedure can synchronize nearly 80% cells at the G1/S transition and minimally impacts the growth and morphology of cells [29, 40] (Appendix Figure S3). Following labeling and fixation, we applied dual-color STORM (Methods) to image the TADs (Figure 1a, green) and the replication initiation sites, which appeared as punctate foci (Figure 1a, purple). The punctate distribution of 10-min pulse labeled foci imaged by STORM in our work were similar with those were previously imaged by other groups with SIM or STORM [23]. The positions of these foci can precisely represent the position of the corresponding replication initiation sites for two reasons. First, as EdU was added immediately after the release of replication arrest, majority of the pulse-labeled RFi were supposed to contain the corresponding replication initiation sites. Second, although the sequence length of the DNA replicated over a 10-min period was quite long (roughly 20 kb [9, 23, 40]), its physical size was only ∼30 nm in diameter, as revealed in the super-resolution images (Figure 1a, Appendix Figure S6c). This diameter is close to the lateral resolution of STORM imaging. As the spatial resolution (directly related with the single-molecule localization precision) sets the minimal apparent size of a target imaged by STORM [33], there would be no difference in the apparent size or position when imaging a 20 kb genomic region and a much smaller sub-region, e.g., the replication initiation site in the region.

**Figure 1.**
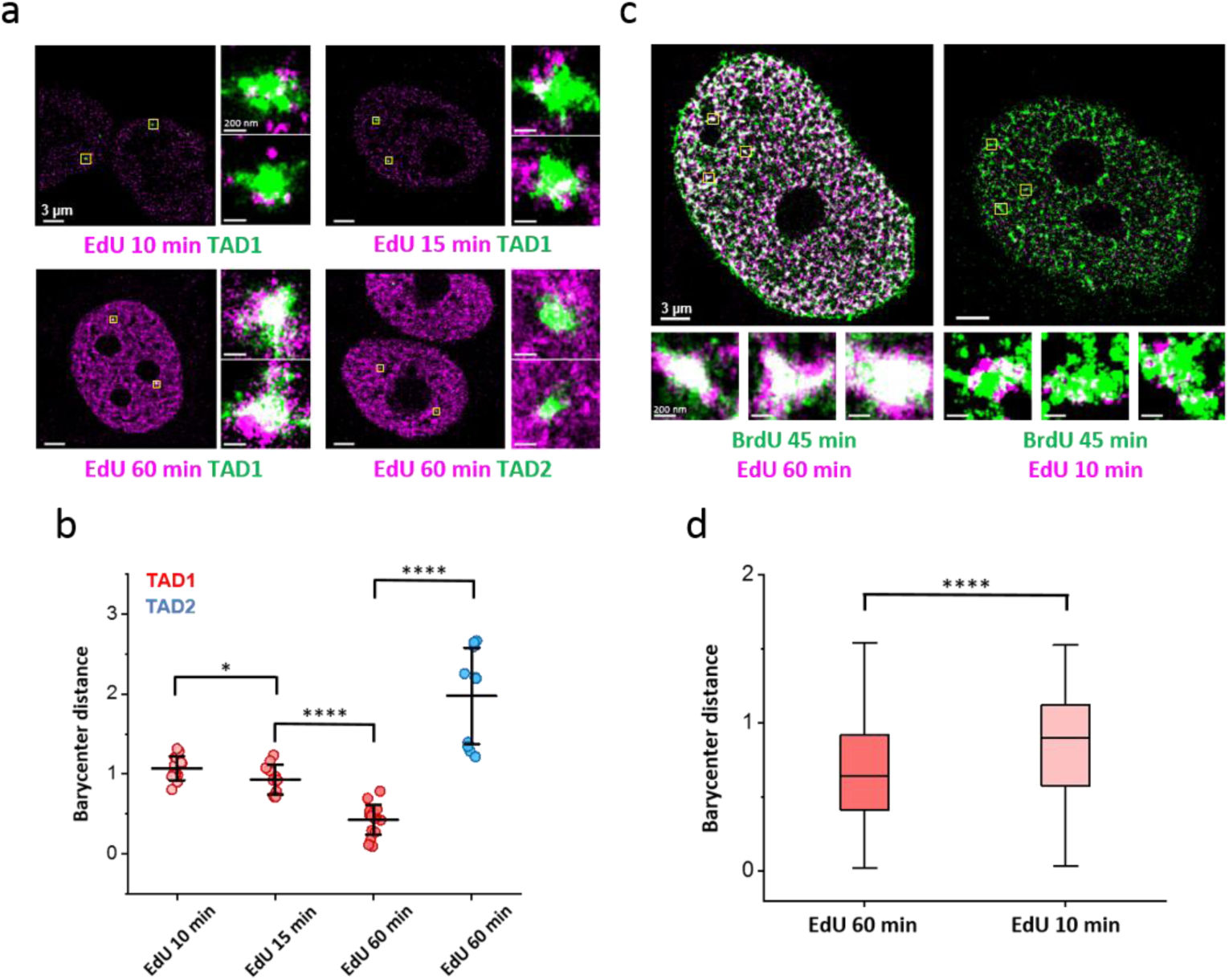
Super-resolution imaging of RFi and TADs in the S phase. **a,** Representative STORM images of TAD1 and TAD2 labeled by Oligopaint probes (green) and RFi labeled metabolically for different durations (purple) (Methods). TAD1 and TAD2 were chosen based on the replication timing profile and Hi-C interaction heatmap of HeLa cells (Appendix Figure S1). TAD1: an early replicating domain (Chr1:16911932-17714928). TAD2: a middle replicating domain (Chr1:17722716-18846245). Metabolic labeling of DNA replication was performed by supplying EdU to the cell upon release into the S phase for 10 min, 15 min, and 60 min (purple). The areas inside the yellow squares are shown at higher magnification to the right of each nucleus. Portions of the two signals that overlap are shown in white. **b,** Barycenter distances between the TAD and its spatially associated RFi (Methods) in **a**. Horizontal lines and error bars represent the mean values ± s.d. of three or more independent biological replicates (n =16 cells). **c,** Representative STORM images of RFi labeled metabolically for different durations in two consecutive cell cycles. Consecutive metabolic labeling of DNA replication was performed by supplying BrdU (green) to the cell upon release into the S phase in the first cell cycle, followed by supplying EdU (purple) to the cell upon release into the S phase in the second cell cycle (for different durations). The areas inside the yellow squares are shown at higher magnification below each nucleus. **d,** Box plot of barycenter distances between BrdU and EdU labeled RFi in **c** (data were pooled from n =10 cells). Center line, median; box limits, 25% and 75% of the entire population; whiskers, observations within 1.5× the interquartile range of the box limits. Significance was analyzed by un-paired two sample parametric t test. ****P < 0.0001, ***P < 0.0005, **P < 0.01, *P < 0.05, N.S.: not significant. 3D results are shown in Fig. S2&S5.

Intriguingly, the replication initiation sites seemed to preferentially localize at the physical boundary of the early replicating TAD1 as shown in the two insets in the upper left panel of Figure 1a, which are close-up view of the two allelic TAD1 and their associated replication initiation sites. We adopted a recently developed robust and unbiased segmentation algorithm, SR-Tesseler [41], to quantitatively analyze TADs and origins in super-resolution images (Methods & Appendix Figure S4). To describe the relative spatial relationship, we defined barycenter distance as the physical distance between the barycenter of a TAD and the barycenter of RFi normalized by the radius of gyration of the TAD (Figure 1b) (Methods). The barycenter is the mass density center of all single molecule localizations in a TAD or RFi. For randomly distributed foci within the TAD, the expected mean barycenter distance is 0.71 (Methods). In contrast, the barycenter distances between the 10-min pulse-labeled RFi and TAD1 were near 1 (Figure 1b), indicating a peripheral distribution of the replication initiation sites relative to TAD1. Importantly, the barycenter distance of RFi labeled for 15 min were closer to the center of TAD1 in comparison with RFi labeled for 10 min (Figure 1b), showing that our method is highly sensitive as a means of detecting the spatial distribution of replication origins in a TAD. Moreover, RFi labeled for 60 min were well overlaid on TAD1 (Figure 1a & Appendix Figure S5b) with barycenter distances near 0.5 (Figure 1b & Appendix Figure S5). As a control, RFi labeled for 1 hour starting at the G1/S boundary did not show obvious overlap with the middle replicating TAD2 (Figure 1a, b), consistent with the fact that TAD2 begins to replicate at approximately 3 hours into the S phase (Appendix Figure S1 & Figure 2a).

**Figure 2.**
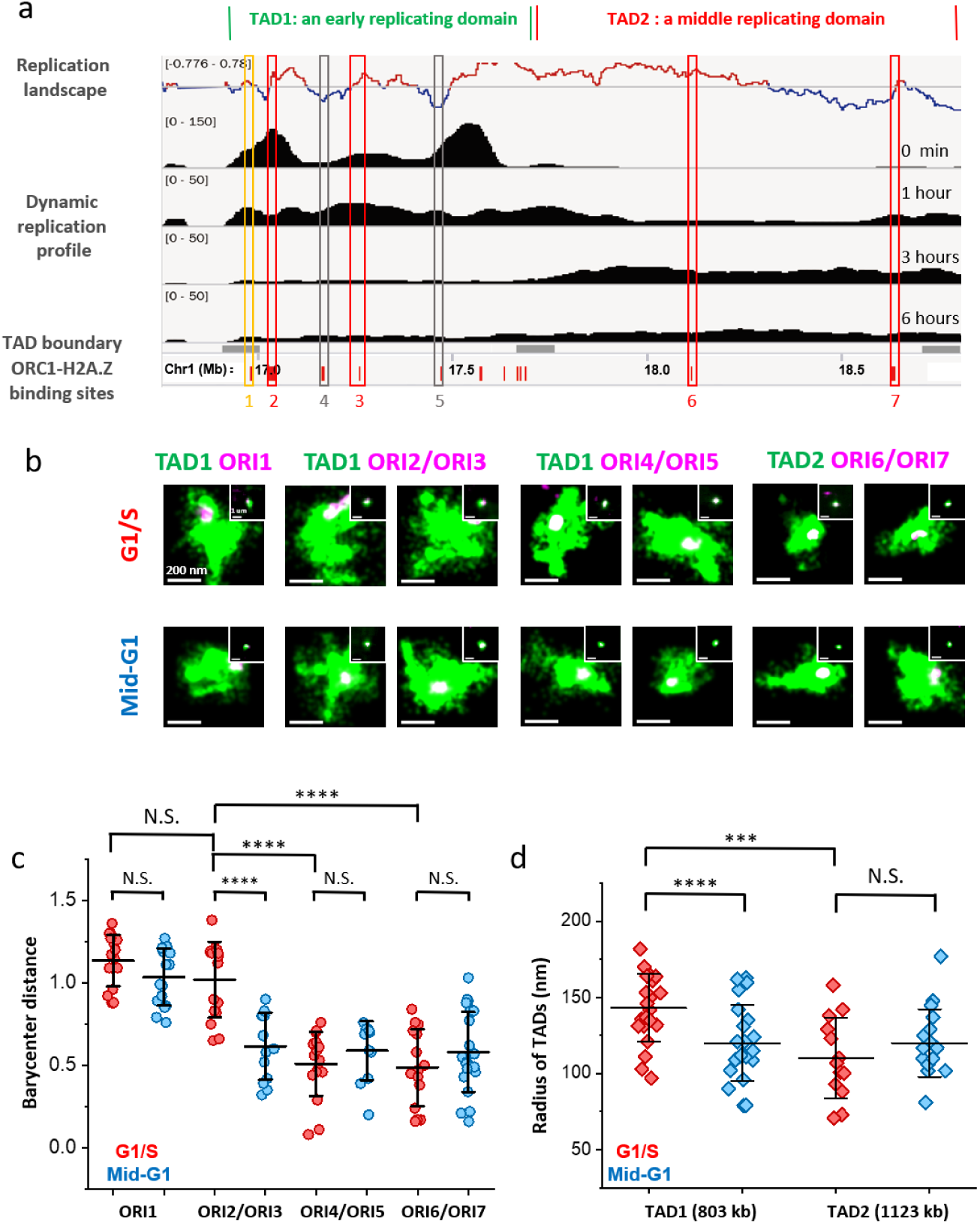
Spatial distribution of replication origins relative to the TADs in the G1 and G1/S phases. **a,** A scheme of replication in TAD1 and TAD2. The top profile represents the replication landscape obtained by OK-seq. (−0.776-0.78) was the threshold of OK-seq [81]. The middle black peaks represent the dynamic replication profile, which was obtained by 10-min BrdU pulse labeling at 0 min, 30 min, 3 hours, and 6 hours into the S phase. (0-50) or (0-150) is the range of normalized BrdU-seq data. The grey bars represent the TAD boundaries in the RDs. The small red bars at the bottom represent the ORC1 and H2A.Z binding sites indicating the potential replication origins. Representative active and dormant replication origins defined by the BrdU-seq data and the OK-seq profile are marked with vertical rectangles. Yellow rectangle: active replication origin (ORI1) at the TAD boundary. Red rectangles: active replication origins in TAD1 (ORI2 and ORI3) and TAD2 (ORI6 and ORI7). Black rectangles: dormant replication origins in TAD1 (ORI4 and ORI5). **b,** Representative STORM images of TADs (green) and their origins (purple) labeled by FISH with oligoprobes in the G1 and G1/S phases. Upper, TADs and origins labeled at the G1/S transition. Lower, TADs and origins labeled at approximately 5 hours into the G1 phase. Portions of the two signals that overlap are shown in white. The corresponding conventional images are shown in the inset. **c,** Barycenter distances between origins and TADs in **b** (n ≥10 cells). **d,** Radii of TAD1 and TAD2 in the G1 or G1/S phase (n ≥10 cells). For lines and statistics in **c** and **d** see the description in the legend of Figure 1. 3D results are shown in Fig. S9.

To further validate the analysis of spatial localization of RFi relative to the TAD, we also applied DBSCAN [42], a density-based spatial clustering algorithm, to extract individual RFi and quantify the spatial localization of RFi in TADs in 2D and 3D STORM images (Appendix Figure S5) (Methods). The spatial relationship of RFi relative to the TAD (Appendix Figure S5a-c) rendered by DBSCAN in 2D or 3D images (Appendix Figure S5a-c) was identical with those obtained by the SR-Tesseler analysis (Figure 1b). We also defined the radial density distribution (RDD), which is the median radial distribution of all single-molecule detections of the RFi in a TAD normalized by the radius of the TAD, to quantify the spatial localization of RFi in TADs in 3D images (Methods). A larger RDD value indicates a more peripheral distribution of RFi in a TAD. The spatial distribution of RFi revealed by the RDD analysis was similar to that obtained by the barycenter distance analysis (Appendix Figure S5c, d). The analyses described above cross validated each other and eliminated the possibility of artifacts introduced by foci identification, inter-foci distance measurement algorithms, or projection of 3D images onto the 2D plane. Therefore, these data demonstrated that multiple replication origins initiate separately at the spatial periphery of TAD1.

Next, to check whether the above findings obtained with TAD1 are generally true for all early replicating TADs, a high-throughput labeling method is needed. Provided that the boundaries of RDs and TADs are precisely aligned [18], we took a metabolic labeling strategy to label all early RDs and their associated replication initiation sites by two rounds of BrdU and EdU pulse labeling at the beginning of the S phase over two consecutive cell cycles, respectively (Figure 1c). EdU was labeled with Alexa647 by click chemistry, whereas BrdU was immunostained with atto-550 [9]. It has been suggested that RFi labeled by thymidine analogs EdU or BrdU for 1 hour generally correspond to RDs [9], as replication of early RDs takes approximately 1 hour [9, 21, 43, 44]. This assumption was validated by large overlapping between FISH-labeled TAD1 and its corresponding 60-min EdU-labeled RFi (Figure 1a lower left, Appendix Figure S5, Appendix Figure S6a, b). Because the spatial density of 60-min metabolically labeled RFi was very high, which impeded the confidence in the algorithm with regard to identification of the boundaries between spatially adjacent RDs, we chose to label RDs using 45-min labeling duration. As shown by the dual-color STORM imaging in Figure 1c, early RDs double-labeled for 45 min and 60 min in two consecutive cell cycles merged very well, in line with the fact that RDs are conserved over cell cycles [9, 21, 25]. RFi labeled for 45 min were slightly smaller than those labeled for 60 min, albeit insignificant (Appendix Figure S6c), supporting the usage of 45-min labeled RFi to represent early RDs. As a control experiment, we showed that the size increase from RFi labeled for 10 min to those labeled for 15 min was successfully detected (Appendix Figure S6c), demonstrating the high detection sensitivity of STORM imaging and also excluding the possibility that the insignificant size difference between RFi labeled for 45 min and 60 min was due to insufficient detection sensitivity. We also checked whether different thymidine analogs, *e.g.* EdU and BrdU, introduced any difference in the size of RFi. The STORM images showed no significant difference between EdU- and BrdU-labeled RFi (Appendix Figure S6d).

We then labeled all early RDs with 45-min BrdU in the first cell cycle followed by labeling of the replication initiation sites with 10-min EdU in the second cell cycle (Figure 1c right). We obtained a global view of the spatial distribution of replication initiation sites relative to early RDs in a cell. Analysis of the dual-color STORM images showed that there were averagely 7 replication initiation sites in each RD, in good agreement with previous estimations [5, 45] as well as the fact that the sizes of a RD and a replicon are about 800 kb and 120 kb, respectively. These analyses thereby benchmarked the technical rigor of labeling and imaging of RDs and associated origins.

We next calculated the barycenter distances between replication initiation sites (EdU 10 min) and RDs (BrdU 45 min) and found that they were significantly larger than those between doubly labeled (EdU 60 min and BrdU 45 min) RDs (Figure 1d). Lastly, a similar spatial pattern was observed when Cy3B or dUTP was used respectively instead of atto-550 or BrdU, thereby eliminating the possibility that the observed pattern could be the consequence of labeling or detection artifacts associated with specific dyes (Appendix Figure S7). Taken together the data of both particular RDs and metabolically-labeled RDs, we conclude the fired replication origins in a RD are spatially separated, which is in direct contrary with the classic model [4] and in line with recent findings discovered by super-resolution imaging [23]. More importantly, these spatially separated replication origins tend to initiate at the periphery of RDs, implicating a role of chromatin domain structure in regulating the efficiency of replication origins.

### Active origins relocate inside-out to the periphery of early replicating TADs in the G1 phase

Only 10–20% of the origins in a TAD are used for DNA replication during each cell cycle, while the rest stay dormant. Given the observation that replication tends to initiate at the periphery of an early replicating TAD (Figure 1 & Appendix Figure S5), we next set to image both active and dormant origins to check whether the spatial distribution of origins in a TAD is related to their replication efficiency. As dormant origins cannot be marked with metabolic pulse labeling, OligoSTORM was applied to label and image the TADs and origins. To obtain the replication efficiency of origins in TAD1 and TAD2, we first measured the dynamic replication profile of HeLa cells using BrdU-seq [46] by 10-min BrdU labeling at 0 min, 1 hour, 3 hours and 6 hours into the S phase (Figure 2a, black peaks). The BrdU-seq profile reveals that TAD1 replicates in the first hour of the S phase, while TAD2 starts to replicate after about 3 hours, which is in line with the replication timing profile (Appendix Figure S1). All potential replication origins in TAD1 and TAD2 were mapped by ORC1 and H2A.Z ChIP-seq [47] of HeLa cells. We aligned the dynamic replication profile and ORC1 binding sites with the previously reported replication landscape of the HeLa cell genome [48] (Figure 2a). Based on the origin efficiency revealed by both the replication landscape and the dynamic replication profile, 3 representative active origins (ORI1, ORI2 and ORI3, marked by yellow and red boxes) and 2 representative dormant origins (ORI4 and ORI5, marked by black boxes) were chosen in TAD1. Two active origins (ORI6 and ORI7, marked by red boxes) were chosen in TAD2 (Additional File Table 1 for detailed information of TADs and origins).

We applied dual-color OligoSTORM to image the TAD and its associated origins at the G1/S transition without releasing aphidicolin arrest (Methods). In order to ensure that the target replication origin was labeled with ample fluorescent signal, Oligopaint probes were designed to target a ∼20 kb genomic zone containing the replication origin (Methods). The results showed that at the G1/S transition, all 3 active origins in TAD1 were located at the spatial periphery of TAD1 (Figure 2b, upper) with large barycenter distances (Figure 2c, red). In contrast to the active origins, the dormant origins (ORI4 and ORI5) tended to locate at the interior of TAD1 (Figure 2b, upper) with barycenter distances much shorter than those of ORI1, ORI2 and ORI3 (Figure 2c, red). Interestingly, the active origins in TAD2 (ORI6 and ORI7), which are not supposed to fire until the middle S phase, tended to locate inside of the domain at the G1/S transition (Figure 2b, upper) with small barycenter distances (Figure. 2c, red). Taken together, these results suggest that the replication efficiency of origins at the G1/S transition is correlated with their physical positions in the TAD.

Eukaryotic DNA replication is tightly orchestrated with the cell cycle. In the canonical two-step activation model [4], licensing of origins occurs with pre-RC formation in the G1 phase followed by origin activation and initiation in the S phase. Recent Hi-C studies have shown that the structure of TADs changes from the G1 phase to the S phase [49]. We wondered how the spatial distribution of origins in a TAD changes accompanying the chromatin structure, which could serve as determinants of selective origin initiation. We thus imaged the TAD and origins in the mid-G1 phase (approximately 5 hours post G1 onset) (Methods), which is after the timing decision point (TDP) when the replication timing program becomes established and TADs reform [50]. Strikingly, we found that the active origins ORI2 and ORI3 were located inside of TAD1 in the mid-G1 phase (Figure 2b, lower) with small barycenter distances (Figure 2c, blue), in sharp contrast to their peripheral localization in TAD1 at the G1/S transition (Figure 2b, upper; Figure 2c, red). On the contrary, dormant origins ORI4 and ORI5 were found to remain inside of TAD1 from the mid-G1 (Figure 2b, lower; Figure 2c, blue) to the G1/S transition (Figure 2b, upper; Figure 2c, red). These observations suggested that active origins undergo an inside-out relocation process in the TAD, possibly along with the chromatin structural re-organization within the TADs that occurs in the G1 phase. Interestingly, unlike ORI2 and ORI3, active origin ORI1 did not relocate but remained at the TAD periphery from the mid-G1 phase to the G1/S transition (Figure 2b, c). We note that, in the sequence space, ORI1 is at the insulation boundary of TAD1 (Figure 2a and Additional File Table 1), and therefore structural re-organization within the TADs would not affect its peripheral localization relative to the TAD. Such correspondence between the sequence boundary and the physical boundary of a TAD was also reported in a recent study [38], thereby again benchmarking the technical rigor of labeling and imaging of TADs and associated origins.

To further investigate the relationship of replication origins and chromatin loops within the TADs, we aligned origins with the sites enriched of CTCF and cohesin genome wide (Methods) (Appendix Figure S8). CTCF and cohesin are the key scaffold protein complexes bound at the anchor sites of the chromatin loops as well as the TAD boundary [51]. In general, compared with random DNA loci, CTCF-cohesin binding sites were enriched with replication origins. Active origins co-localized better with CTCF-cohesin binding sites than dormant origins. In addition, active origins located at TAD boundaries (Methods) were of higher replication efficiency than those located inside the TADs. The sequencing results again emphasized the relationship of replication efficiency with chromatin organization within the TADs.

In addition to the structural re-organization within the TADs, we found that the physical size of TAD1 also became larger at the G1/S transition in comparison with its size in the G1 phase (Figure 2d), while this change was not detected for TAD2. Note that the volume increase was not the result of DNA replication, as the cells were arrested at the G1/S transition, suggesting that the chromatin of TAD1 undergoes de-compaction in the G1 phase, which is in line with the results of Hi-C analysis showing that intra-domain chromatin interactions decrease in the G1 phase [49]. Analysis of 3D STORM images led to the same findings (Appendix Figure S9), which again eliminated the possibility of artifacts introduced by projection of 3D images onto the 2D plane. Taken together, these data revealed that the structural re-organization within the TADs and decompaction in the G1 phase facilitate the relocation of active origins from the TAD interior to the periphery, supporting the observation that DNA replication initiates at the periphery of TADs in the beginning of S phase (Figure 1).

### Distinct spatial localization of active and dormant origins at the G1/S transition is correlated with chromatin loops and dependent on transcription

Next, we set to explore the factors that are responsible for differential origin localization in the TAD. We first examined the effects of CTCF and cohesin, the key scaffold protein complexes responsible for loop formation in a TAD [51]. Upon down-regulation of CTCF or cohesin using RNAi (Figure 3a, insets), we found that the active origins (ORI2 and ORI3) was not relocated to the TAD periphery at the G1/S transition in both 2D and 3D images (Figure 3a & Appendix Figure S10a). More importantly, the barycenter distances of either active origins or dormant origins relative to TAD1 became similar with that of randomly distributed foci (about 0.7) (Figure 3b & Appendix Figure S10b), indicating disappearance of differential origin distribution. Such effect was likely due to the scrambling of chromatin structure within the TADs upon loss of CTCF or cohesin, which is in line with the Hi-C data that depletion of either cohesin or CTCF eliminates loops [52, 53]. These results suggested that the relocation of replication origins in the G1 phase is dependent on chromatin looping mediated by CTCF and cohesin in the TAD.

**Figure 3.**
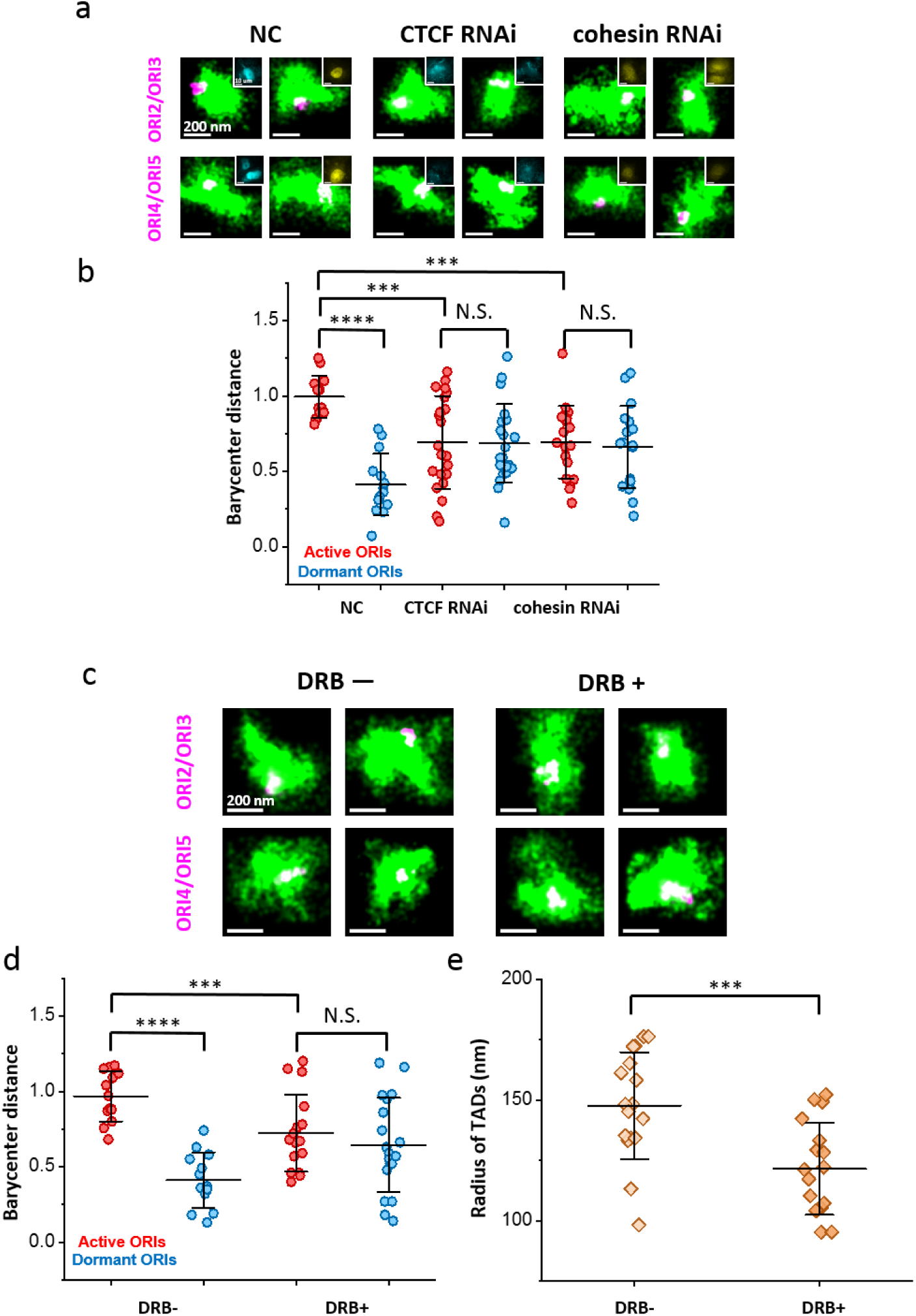
The spatial distribution of replication origins within the TADs at the G1/S transition is dependent on CTCF, cohesin and transcription. **a,** Representative STORM images of origins (purple) in TAD1 (green) after treatment of cells with the indicated siRNAs. Conventional images indicate the concentration of CTCF (cyan) or cohesin (yellow) in the nucleus. **b,** Barycenter distances between active or dormant origins in TAD1 after treatment of cells with the indicated siRNAs. Portions of the two signals that overlap are shown in white. **c,** Representative STORM images of origins (purple) in TAD1 (green). Left: no DRB. Right: with DRB. **d,** Barycenter distances between active/dormant origins and TAD1 with or without DRB treatment. **e,** Radius of TAD1 treated with or without DRB. For lines and statistics in **b**, **d**, and **e** see the description in the legend of Figure 1 (n ≥10 cells). 3D results are shown in Fig. S10.

Previous studies have shown that in the G1 phase, transcription activity is generally high in early RDs and active origins abut actively transcribed genes [48, 54]. Indeed, expression of genes associated with active replication origins (ORI1, ORI2, ORI3) in TAD1 is higher than that for dormant replication origins (ORI4, ORI5) (Appendix Figure S11). Transcription has also been found to fundamentally influence chromatin structures at different levels and through various mechanisms, including nucleosome disassembly, enhancer-promoter loop formation, transcript cis-interaction, CTCF and cohesin displacement, gene relocation and transcription factory formation [55–57]. Moreover, in a recent study, Gilbert and his colleagues identified cis-acting elements, namely early replicating control elements (ERCEs), which regulate the replication timing and the structure of TADs [58]. Importantly, ERCEs have properties of enhancers or promoters, implicating a fundamental role of transcription in orchestrating genome replication and chromatin architecture. Therefore, given that the origin relocation takes place in the G1 phase, we next examined whether transcription in the G1 phase contributes to chromatin structural re-organization within the TADs. To do so, we treated cells with transcription elongation inhibitor 5,6-dichloro-1-β-D-ribofuranosyl-benzimidazole (DRB) [59] from the mid-G1 phase to the G1/S transition, after which we labeled TAD1 and its replication origins using Oligopaint probes. Interestingly, upon transcription inhibition by DRB treatment, active origins ORI2 and ORI3 were no longer found to relocate to the periphery of the TAD at the G1/S transition in both 2D and 3D images (Figure 3c & Appendix Figure S10c) and had barycenter distances similar to those of dormant origins (Figure 3d & Appendix Figure S10d). Moreover, the radius of TAD1 at the G1/S transition in DRB-treated cells was found to be smaller than that in normal cells (Figure 3e & Appendix Figure S10e) and similar with that observed in the G1 phase (Figure 2d & Appendix Figure S9c). This observation suggested that transcription de-compacts the chromatin structure of TADs. Taken together, these results demonstrated that transcription-dependent chromatin structural re-organization within the TADs exposes a subset of origins to the physical boundary of a TAD, which are preferentially used for replication initiation.

### Replication elongation factor PCNA surrounds TADs both in the G1 and G1/S phases

To answer why origins located on the physical boundary of a TAD are preferentially used for DNA replication, we examined the spatial distribution of replication machinery relative to individual TADs at the G1 phase and G1/S phases by imaging proliferating cell nuclear antigen (PCNA) [60]. As a control, we also monitored the distribution of minichromosome maintenance complex component 2 (MCM2) and CTCF, respectively. Provided that metabolically labeled RDs merge well with FISH-labeled TADs in the early S phase (Figure 1, Appendix Figure S5 and Appendix Figure S6), to label early replicating TADs and protein factors in the same cell, TADs were first labeled by supplying cells with EdU for 45 mins immediately after release of aphidicolin arrest at the beginning of the S phase; in the next cell cycle, the cells were fixed and immunostained at either the mid-G1 or G1/S phase. The EdU-labeled TADs became larger from the mid-G1 phase to the G1/S transition (Appendix Figure S12), in line with the observation of FISH-labeled TADs (Fig. 2d). As a scaffold factor of TADs, CTCF formed large foci (Figure 4a) and neither their distribution relative to the TADs (Figure 4d) nor their sizes (Figure 4e) was found to change in the G1 phase. Interestingly, despite the constant sizes of the CTCF foci, both the single-molecule detection counts (Appendix Figure S13a) and the molecule density (Appendix Figure S13b) in the CTCF foci were reduced from the mid-G1 phase to the G1/S transition, suggesting that CTCF dissociated from DNA during the transcription-dependent chromatin structural re-organization process. This STORM-based finding is consistent with a previous single-molecule study showing that binding of CTCF to chromatin decreases from the G1 phase to the S phase [61], as well as a Hi-C study showing that transcription elongation can disrupt CTCF-anchored chromatin loops [57].

**Figure 4.**
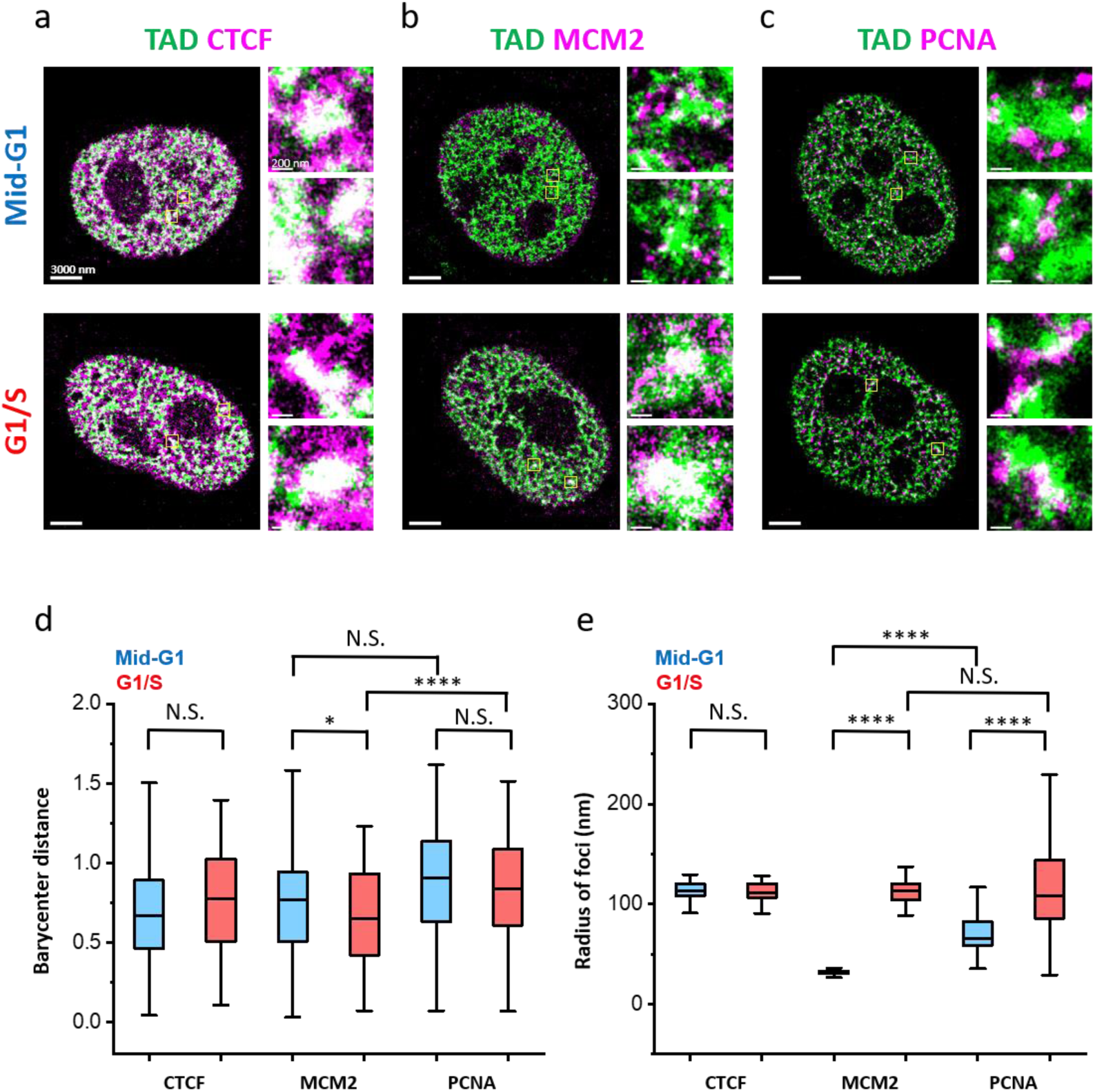
Spatial distributions of CTCF, MCM2 and PCNA relative to the early replicating TADs in the G1 and G1/S phases. **a-c,** Representative STORM images of CTCF, MCM2 and PCNA labeled by immunostaining (purple) and metabolically labeled TADs (green). Cells were fixed and labeled in the mid-G1 phase (upper) or G1/S phase (lower). The areas inside the yellow squares are shown at higher magnification next to each nucleus. Portions of the two signals that overlap are shown in white. **d,** Barycenter distances between CTCF, MCM2 or PCNA with the TADs in the mid-G1 phase or G1/S phase. **e,** Radii of CTCF, MCM2 or PCNA foci in the mid-G1 phase or G1/S phase. For lines and statistics in **d** and **e** see the description in the legend of Figure 1 (n =10 cells).

In contrast to CTCF, MCM2 and PCNA showed drastically different patterns. In the G1 phase, MCM2 formed small clusters that distributed relatively around the TADs (Figure 4b, d). Intriguingly, at the G1/S transition, MCM2 foci became significantly larger (Figure 4b, e) and distributed more toward the center of the TADs (Figure 4b, d). Quantitative analyses of the foci showed that, while the single-molecule detection counts in the MCM2 foci increased from the mid-G1 phase to the G1/S transition (Appendix Figure S13a), the molecule density decreased (Appendix Figure S13b). This observation suggested that MCM2 gradually became associated with chromatin in the G1 phase, in line with the results by Gilbert and his colleagues [62]. As a replication elongation factor of the initiation complex, PCNA binds a subset of origins with the pre-IC complex and recruits DNA polymerases. In the G1 phase, we found that PCNA formed small clusters around the TADs (Figure 4c, d). However, unlike MCM2, the PCNA foci remained surrounding the TADs at the beginning of the S phase (Figure 4c, d). This spatial distribution of PCNA provides a plausible reason why origins at the TAD periphery get preferentially initiated. Moreover, from the mid-G1 phase to the G1/S transition, the size of the PCNA foci was nearly doubled (Figure 4c, e) with both the single molecule detection counts and the molecule density in the foci increased dramatically (Appendix Figure S13a & b). These data suggested that PCNA was gradually recruited to chromatin DNA, consistent with the previous reports that PCNA clusters are much more visible by live cell imaging in the S phase in comparison with the G1 phase [63–66].

## DISCUSSION

Here, we unveiled a new mechanism for replication origin selection by directly visualizing individual TADs and the spatial distribution and dynamics of replication origins in the TADs using super-resolution imaging. We first found that replication initiation generally takes place separately at the spatial boundary of the TAD (Figure 1 and Appendix Figure S5). Next, we discovered that origins undergo relocalization along with the structural re-organization within the TAD in the G1 phase, and the origins that either relocate to (*e.g.* ORI2 and ORI3) or remain at (*e.g.* ORI1) the spatial boundary of the TAD are of higher replication efficiency (Figure 2 and Appendix Figure S9). Importantly, we found that chromatin structural re-organization within the TADs is driven by disruption of chromatin loops during transcription elongation [57] in the G1 phase (Figure 3 and Appendix Figure S10). Lastly, we observed that the major replication machinery protein PCNA, which was previously found to be immobile in the S phase [63–66], remains surrounding the TADs from the mid-G1 phase to the S phase and provides the origins exposed at the spatial boundary of a TAD with a better chance of accessing the replication machinery.

### The “Chromatin Re-organization Induced Selective Initiation” (CRISI) Model

Based on our results, we propose a “Chromatin Re-organization Induced Selective Initiation” (CRISI) model (Figure 5) for replication origin selection. The CRISI model suggests that the spatial localization of an origin in a TAD determines its replication efficiency. Dynamically, in the early-to-mid G1 phase, all potential origins distribute homogeneously in the TAD (Figure. 5a). Upon the onset of transcription, the chromatin loops in the TAD are de-compacted and some loop anchors are disrupted, leading to a subset of origins relocalizing from the inside of the TAD to the periphery (Figure 5b, c). Meanwhile, PCNA forms clusters that remain around the TAD from the mid-G1 phase to the G1/S transition. The peripherally and separately located origins are more accessible to the surrounding PCNA clusters and thus become active origins (Figure 5c). The distribution of active origins in TADs in our CRISI model is in contrary with the classic Rosette model, which proposes that the active origins cluster and co-fire in the chromatin domain [4].

**Figure 5.**
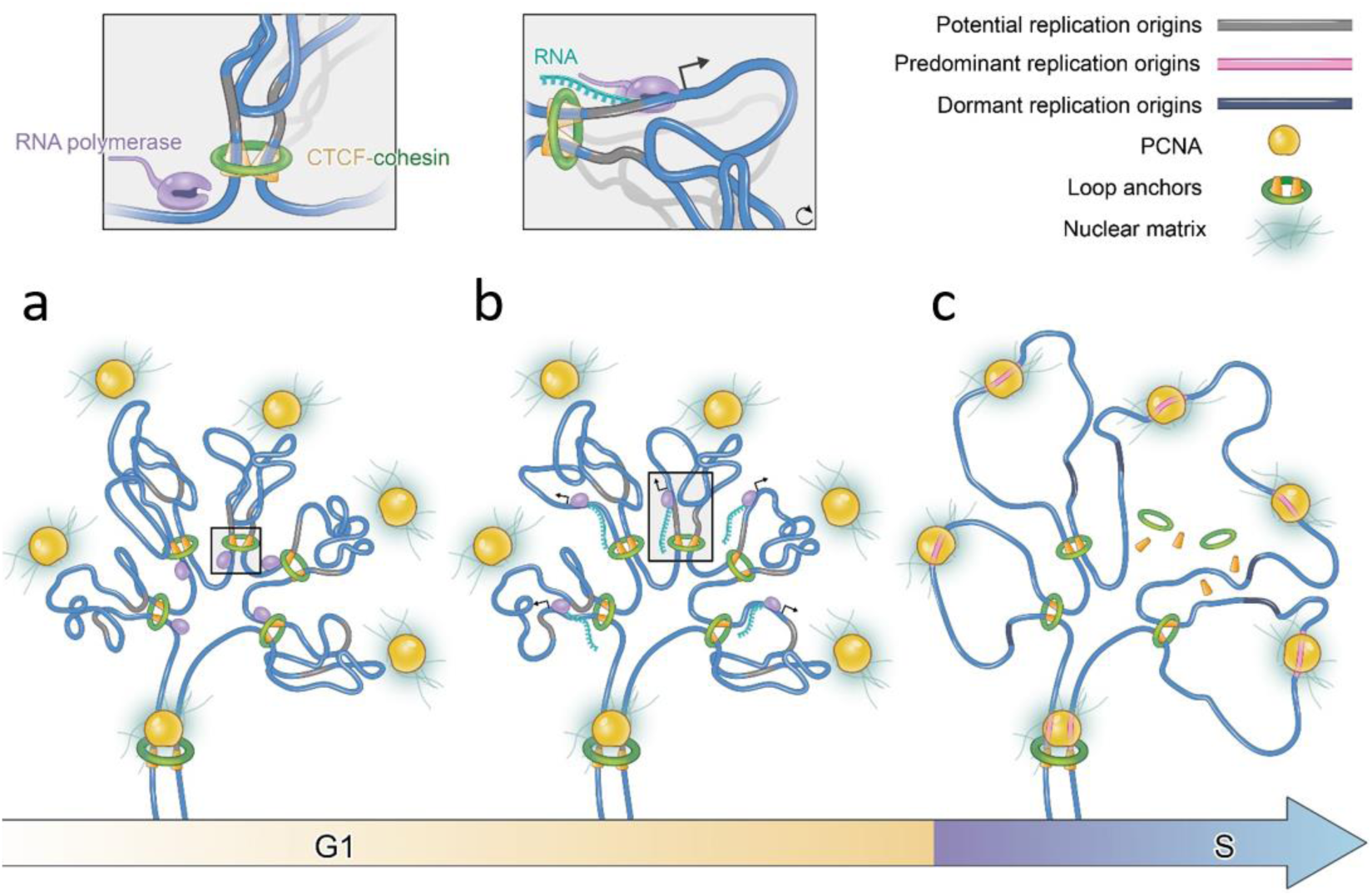
“Chromatin Re-organization Induced Selective Initiation” (CRISI) model for selective initiation of DNA replication origins. **a,** In the early G1 phase, the spatial distributions of potential replication origins (grey ribbons) are relatively even in the TAD. The TAD comprises several chromatin loops (blue) organized by CTCF and cohesin at the loop anchors (green rings). PCNA clusters (yellow balls) surrounding the TAD are bound to the nuclear matrix (hazed light blue straws). **b-c,** With transcription proceeding, the chromatin loops undergo structural re-organization along with chromatin domain decompaction in the G1 phase, exposing a subset of the origins to the periphery of the TAD (pink ribbons). Note that the origin at the sequence boundary of the TAD remains at the TAD periphery in the G1 phase. These peripheral origins are more accessible to the surrounding PCNA clusters and thus become active origins for the initiation of DNA replication at the periphery of the TAD. The areas inside the black squares in **a** and **b** are shown at higher magnification above.

Recently, based on the observation that nearly all replicons are spatially separated at the beginning of the S phase, Cardoso and his colleagues also questioned the Rosette model [23]. They proposed a stochastic, proximity-induced replication initiation model, describing induced domino-like origin activation that may lead to the temporal grouping of active replicons within a chromatin fibre [28]. Nevertheless, as the replication origins were not imaged along with the chromatin domains, how the origins are spatially organized in the chromatin domain and whether the distribution can differentiate the origin efficiency were not known. In the current work, we realized the first direct visualization and quantification of the relative localization and organization of replication origins within individual TADs. Given that a TAD is typically ∼200 nm in radius (Figure 2d and Appendix Figure S9c) and the difference in barycenter distances between active and dormant origins is less than 100 nm (Figure 2c and Appendix Figure S9b), such quantification would require simultaneous imaging of both individual TADs and the associated replication origins with nanometer spatial resolution. Therefore, 3D STORM with 20 nm lateral and 50 nm axial resolution would be more suitable for such analyses.

While aphidicolin is commonly used in the investigation of DNA replication [8–10], it is concerned that such replication stress may stimulate the engagement of dormant origins in the activated replication domains [6]. In our study, to avoid selection of abnormally activated dormant origins, we combined the dynamic replication profile measured by BrdU-seq under aphidicolin treatment and the replication landscape measured by OK-seq without aphidicolin treatment. With this strategy, we were able to select dormant replication origins (ORI4 and ORI5), which were not affected by aphidicolin treatment, for FISH labeling in the TADs. For metabolic labeling of replication initiation sites with aphidicolin treatment, although some dormant origins have the chance to be fired under the replication stress (Figure 1), it does not affect the conclusion that replication starts at the periphery of the chromatin domains.

Regarding the constrained mobility of PCNA clusters in the nucleus, we speculate three possible mechanisms that are not mutually exclusive. Firstly, PCNA and the replisomes are giant complexes binding DNA with low diffusive mobility. Secondly, PCNA and the replisomes may be attached to the nuclear matrix (Figure 5c), which is supported by an immunoelectron microscopy study showing that DNA polymerase ɑ, PCNA, and nascent DNA are colocalized in nucleoskeleton bodies [67]. Thirdly, the proteins comprising replisome complexes might form liquid condensates that phase-separate from TADs [68]. These possibilities would be interesting subjects for future studies.

That the chromatin structural dynamics within the TADs make origins accessible to immobile PCNA clusters provides an interesting viewpoint to understand protein-DNA interactions in the nucleus, which are commonly considered to be based on diffusive search of proteins such as transcription factors on chromatin DNA [69, 70]. Such mechanisms may be involved in various nuclear functions. For example, during DNA damage repair, ATM is restricted at the double strand breaking (DSB) site while phosphorylation of H2AX by ATM spreads over a domain [71]. The discrepancy between the distribution of the kinase (ATM) and its product (γH2AX) can be explained by the local movements of the chromatin fiber inside the TAD which bring distant nucleosomes to spatial proximity of ATM [71]. In the future, other imaging methods such as sequential imaging approach (Hi-M) may be combined with oligoSTORM to further investigate chromosome organization and functions in single nuclei [72].

### Replication, transcription, and chromatin structure

It has been known for many years that transcription is profoundly related to replication [73, 74]. However, while transcription is known to be highly correlated with the replication timing of TADs [58, 75], it is unclear whether transcription regulates origin selection within individual TADs. Intriguingly, although the genetic and epigenetic signatures of active origins mapped by various methods seem quite different and hierarchical [2], they are mostly markers of active transcription and interdependent in the context of transcription. Transcription has been reported to change chromatin structure at different levels. Our imaging data reveal that the transcription activities can displace CTCF from the TADs (Appendix Figure S13) and de-compact the chromatin domain (Figure 2d&3e), consistent with the previously reported Hi-C data [49, 57]. These effects, together with other transcription-induced changes in nucleosomes, chromatin fibers and enhancer-promoter loops, re-organize the chromatin structures within the TADs to relocate a subset of origins to the TAD periphery and consequently, these origins possess higher replication efficiency for being more accessible to the peri-TAD PCNA clusters. The transcription-dependent CRISI model predicts that enhancing transcription activity should increase the selectivity of replication origins, while repressing transcription should cause the opposite effect. Indeed, two recent studies of replication initiation in particular genes showed that enhanced transcription leads to more selective initiation of origins [54, 76] while transcription inhibition causes more origins to be used for replication [77]. However, while our results explain why a subset of origins is preferentially activated in a TAD, the envision that how specific origins are relocated to the TAD periphery by transcription activity is unclear. Further investigations using CRISPRi [78] to interfere with the transcription of specific genes and observation of changes in the replication efficiency of origins associated with these genes would illuminate the detailed mechanisms underlying the non-random, yet flexible, nature of replication origin selection.

Encountering between the transcription and replication machineries is a major intrinsic source of genome instability [73, 79]. Therefore, how cells prevent or resolve the transcript-replication conflicts has been an important question. One major mechanism is to temporally separate transcription and replication for the same genomic regions [80]. Our data suggested that the replication machineries are confined around the TAD and spatially separated from the transcription machineries, which mainly function within the TAD. Therefore, our work provides a new mechanism for cells to avoid the conflicts between replication and transcription based on spatial/topological separation.

In summary, the CRISI model demonstrates important coordination among DNA replication, transcription, and chromatin structure, which reconciles the discrepancy of different signatures for origin efficiency. Lastly, our work also provides new insights into how 3D genome structural dynamics, particularly the intra-TAD physical structures, may regulate other nuclear processes on chromatin templates such as DNA repair, adding a new layer of understanding to chromatin structure and functions.

**Supplementary Information** is available in the online version of the paper.

## Acknowledgements

The authors thank Dr. Wei Guo (University of Pennsylvania) for the HeLa-S3 cell line, Florian Levet (University of Bordeaux) for SR-Tesseler analysis and Ruifeng Li (Peking University) for the Hi-C interaction map. This work was supported by grants from the National Key R&D Program of China (No. 2017YFA0505300), the National Natural Science Foundation of China (21573013), and the National Science Fund for Distinguished Young Scholars (21825401 to Y.S.). Grants for G.L. from the Ministry of Science and Technology of China (2017YFA0504200) and the Chinese Academy of Sciences (CAS) Strategic Priority Research Program (XDB19040202).

## Author Contributions

Y.L., B.X. and Y.S. conceived the project and designed the experiments. Y.L. performed the probe synthesis, sample preparation, imaging and data analysis. B.X. performed coding for the data analysis algorithms. H.L. performed the sequencing experiments and L.Z. performed the sequencing data analysis. Y.L. and Y.S. wrote the manuscript. The other authors took part in experiments and data analysis with Y.L.

## Author Information

Reprints and permissions information is available at. The authors declare no competing financial interests. Readers are welcome to comment on the online version of the paper. Correspondence and requests for materials should be addressed to Y.S.

## Figures and Figure Legends

**Appendix Figure S1.**
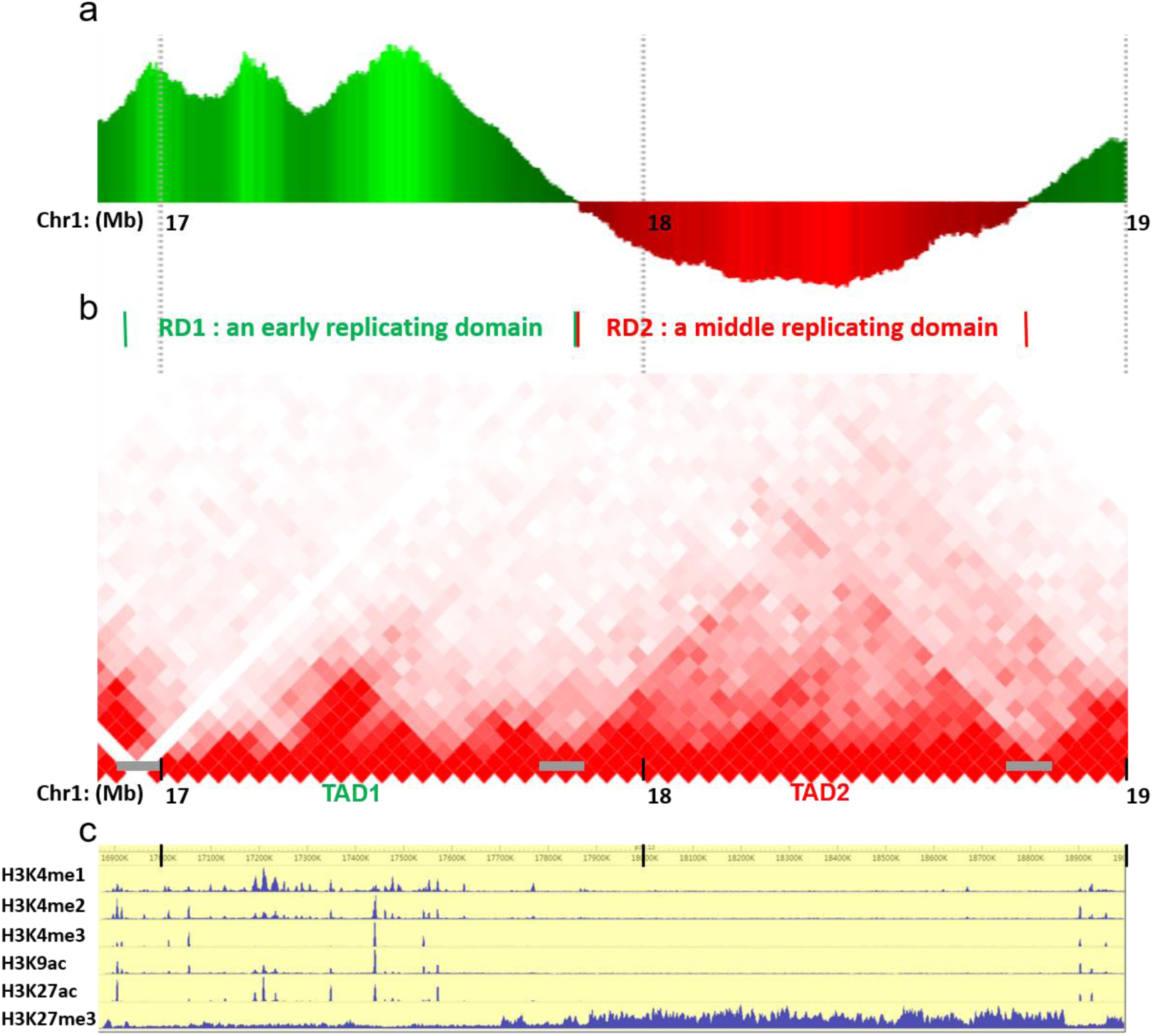
Identification and histone modifications of two TADs from the replication timing profile and Hi-C interaction heatmap of HeLa cells. **a-b,** Depiction of the replication timing profile (green peaks for early RDs and red peaks for middle or late RDs) and Hi-C interaction heatmap in the same genomic region of chromosome 1. The replication timing profile was obtained from the Replication Domain Genome Browser of the Gilbert lab (https://www2.replicationdomain.com/genome_browser). The Hi-C interaction heatmap was obtained from ENCSR693GXU. Grey bars: TAD boundaries. (Methods) Two TADs were selected. TAD1: an early replicating domain (Chr1:16911932-17714928). TAD2: a middle replicating domain (Chr1:17722716-18846245). **c,** Profiles of histone modifications are from public data hubs (ENCODE data portal) of WashU Epigenome Browser.

**Appendix Figure S2.**
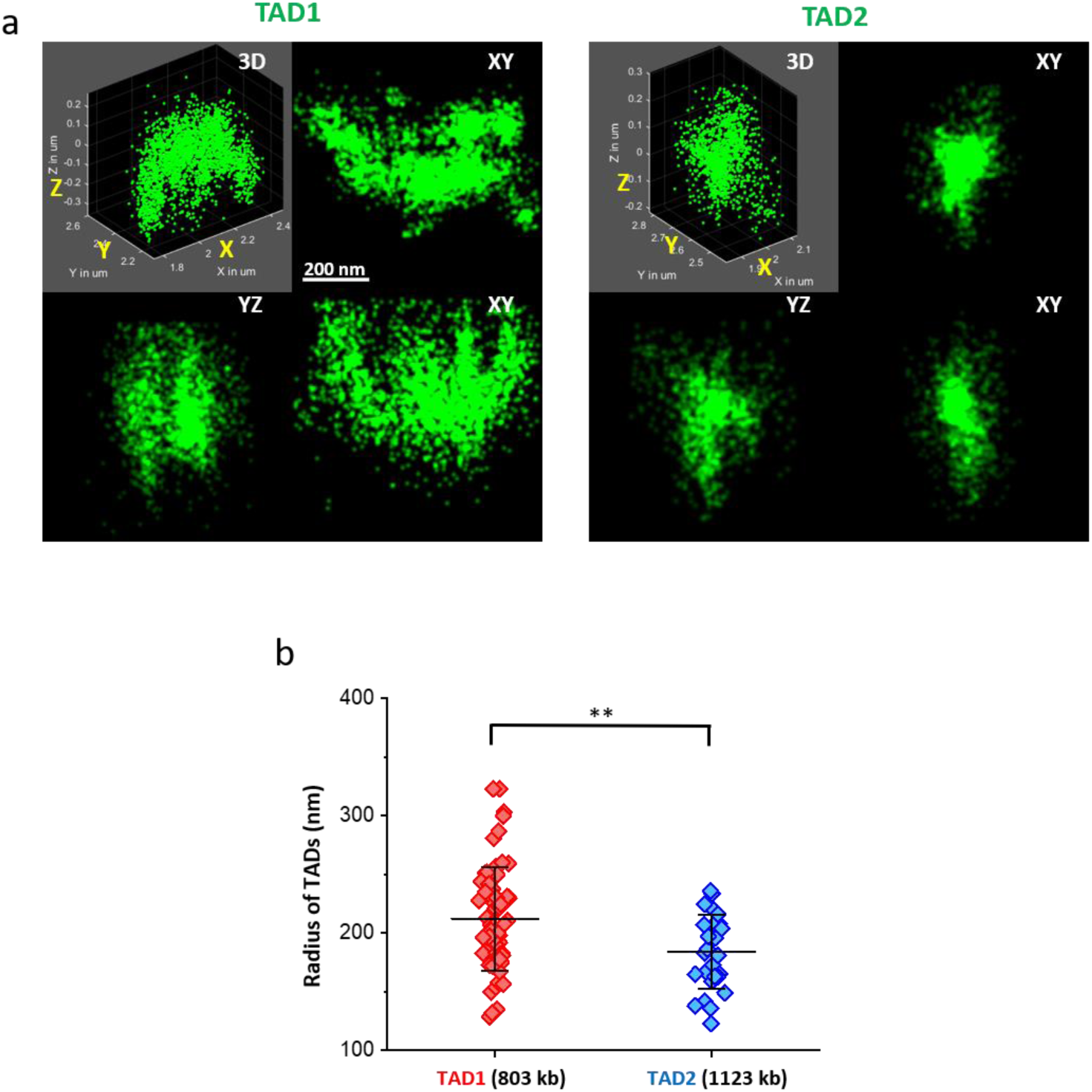
3D visualization and radii of TAD1 and TAD2. The definitions and labeling procedures of TADs and origins are identical with those in Figure 2. **a**, Representative 3D STORM images of TADs. One 3D presentation with 3 projected images. **b,** 3D radius of gyration of TAD1 and TAD2. For lines and statistics in **b** see the description in the legend of Figure 1 (n ≥20 cells).

**Appendix Figure S3.**
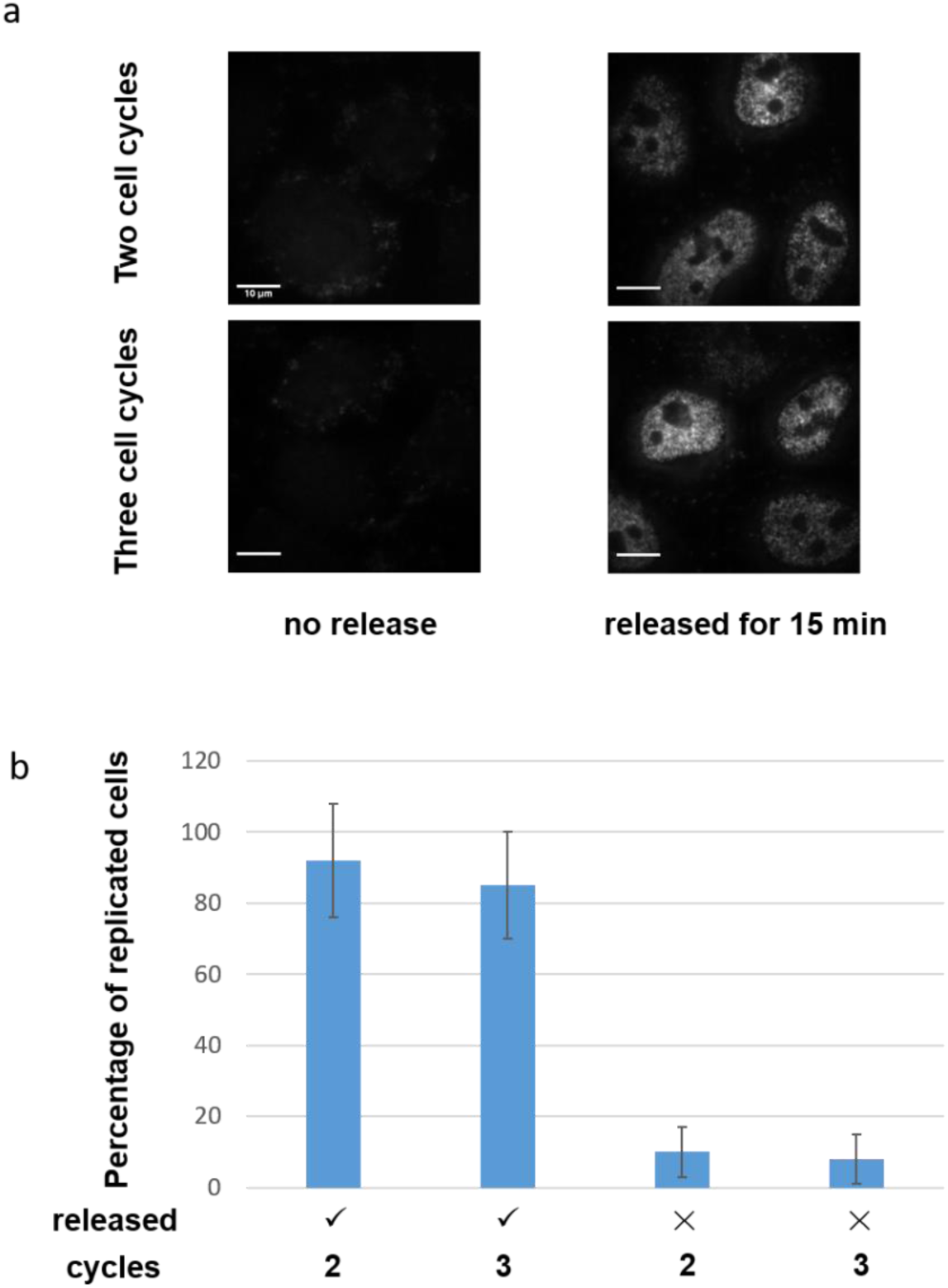
Quantification of cell synchronization by EdU labeling. To make sure the cells were successfully synchronized and the synchronization procedure minimally impacts the growth and morphology of cells, we imaged and analyzed the cells synchronized for two or three cell cycles. **a**, after two (upper) or three (lower) cycles of synchronization, cells were synchronized to the G1/S transition. EdU were added when the synchronized cells were released (right) for 15 min or were not released (left). b, percentage of the replicated cells in **a**. Replicated cells were defined by the three folds of the fluorescence of the nucleus to the background (3 replicates, 200 cells for each group). EdU labeling showed more than 80% cells entering the S phase, similar with the previous work [1].

**Appendix Figure S4.**
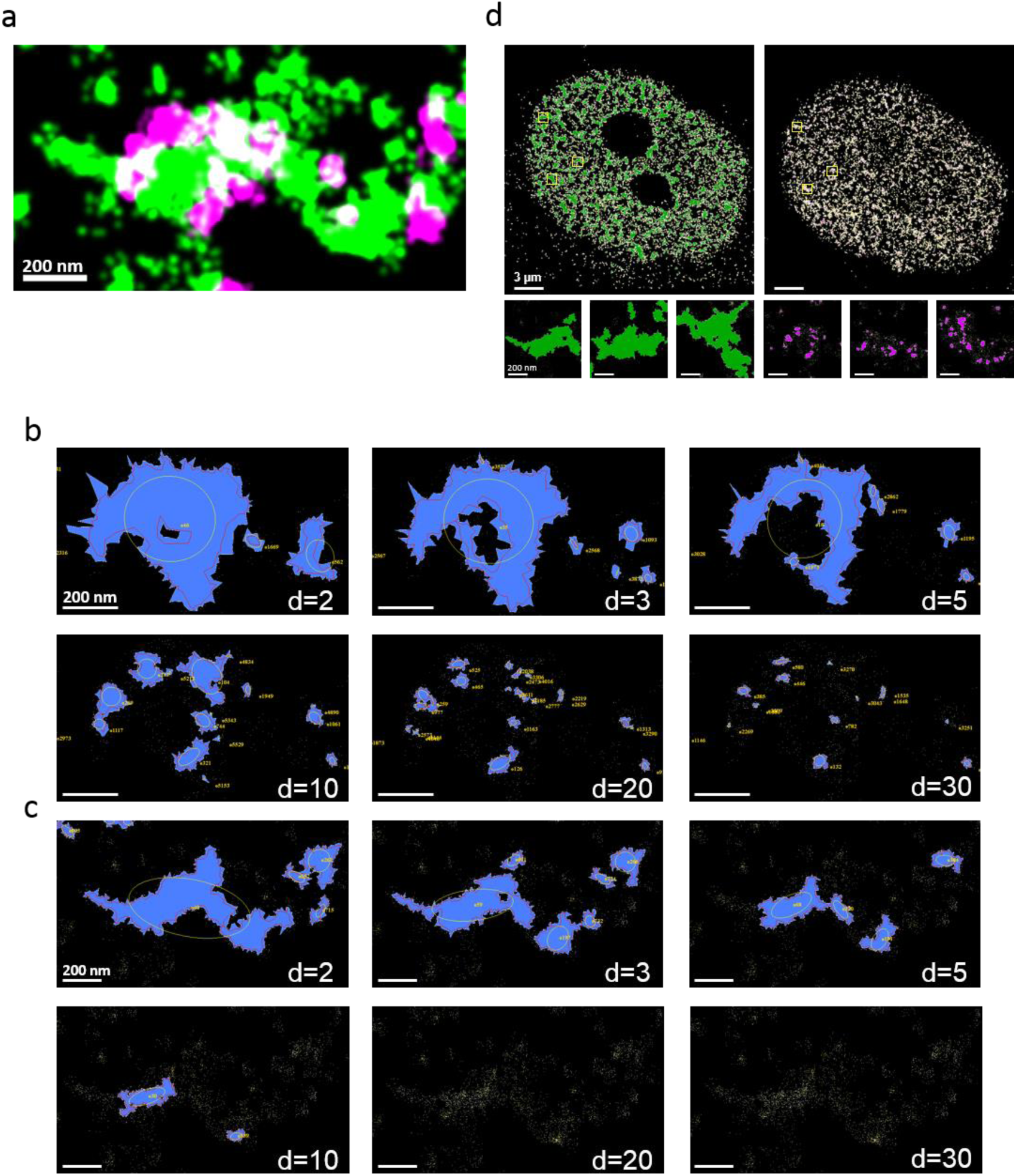
Quantification of the density factor in SR-Tesseler analysis. **a,** Dual-color STORM imaging of TADs (also early replicating domains, green) and origins (purple) as in Fig. 1c. **b** and **c,** Analysis of TADs and origins by SR-Tesseler with density factors from 2 to 30. **b**, Origins close to each other in a TAD cannot be separated when the density factor is set from 2 to 10. Origins are too small or even dismissed when the density factor is set to 30. When the density factor was set to 20, approximately 5,000 origins were clearly defined at the beginning of the S phase in one cell, similar with a previous report [2]. **c**, TADs close to each other cannot be separated when the density factor is set to 2. TADs are too small or even dismissed when the density factor is set from 5 to 30. When the density factor was set to 3, approximately 700 TADs were clearly identified, similar with a previous report[3]. **d**, TADs (green) and origins (purple) identified by SR-Tesseler from the STORM images using a density factor of 3 for TADs and a density factor of 20 for origins. The areas inside the yellow squares are shown at higher magnification below each nucleus.

**Appendix Figure S5.**
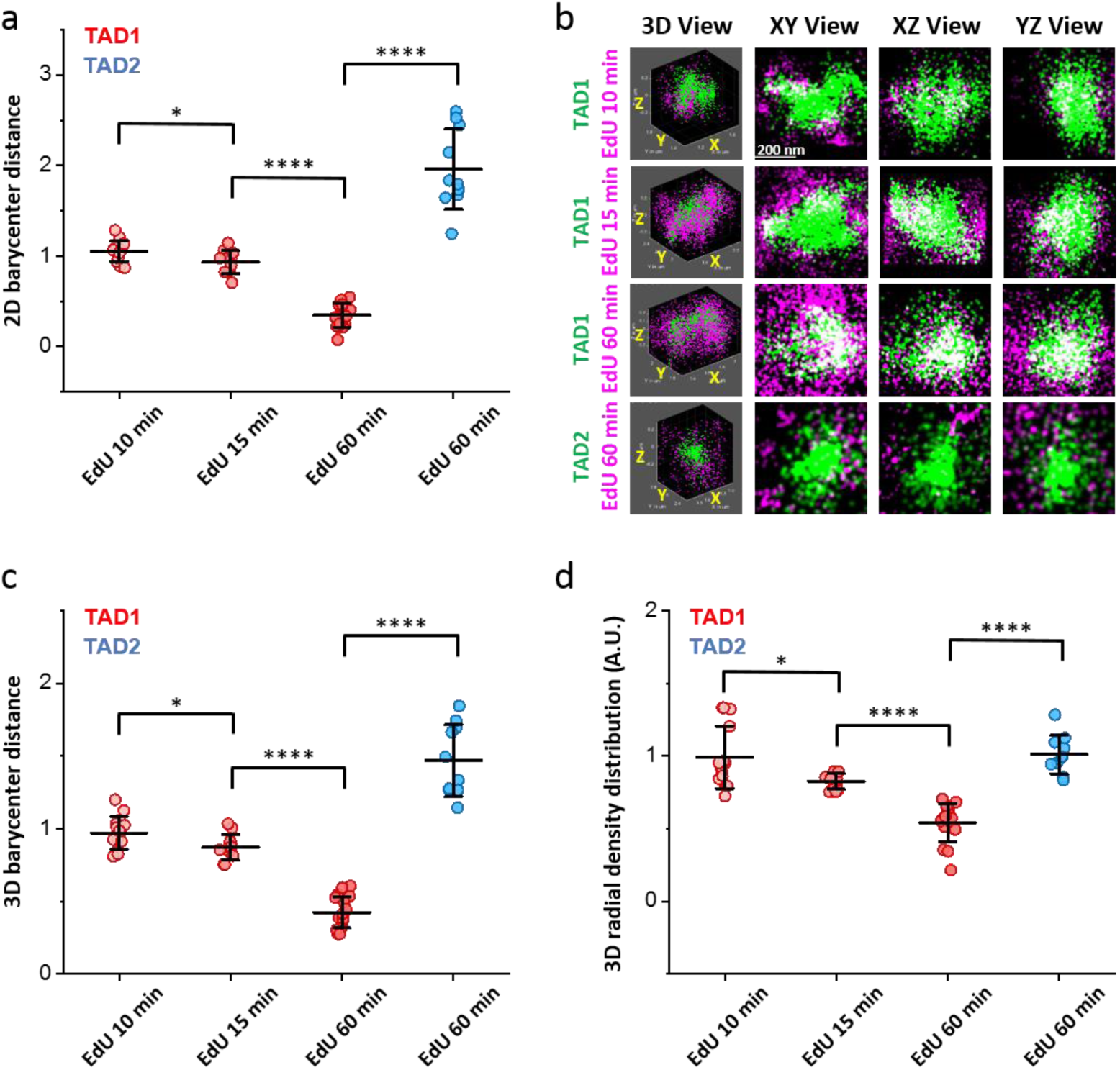
Replication patterns of TADs in the S phase as determined by DBSCAN. **a,** 2D barycenter distances between the TADs and their spatially associated RFi as determined by DBSCAN (Methods) in Figure 1a. **b,** Representative 3D STORM images of TAD1 and TAD2 labeled by Oligopaint probes (green) and RFi labeled metabolically for different durations (purple). Metabolic labeling of DNA replication was performed by supplying EdU to the cells upon release into the S phase for 10 min, 15 min, and 60 min (purple). **c,** 3D barycenter distances between the TADs and their spatially associated RFi as determined by DBSCAN in **b**. **d,** Radial density distribution of the RFi in TADs as determined by DBSCAN (Methods) in **b**. For lines and statistics in **a**, **c**, and **d** see the description in the legend of Figure 1 (n ≥10 cells).

**Appendix Figure S6.**
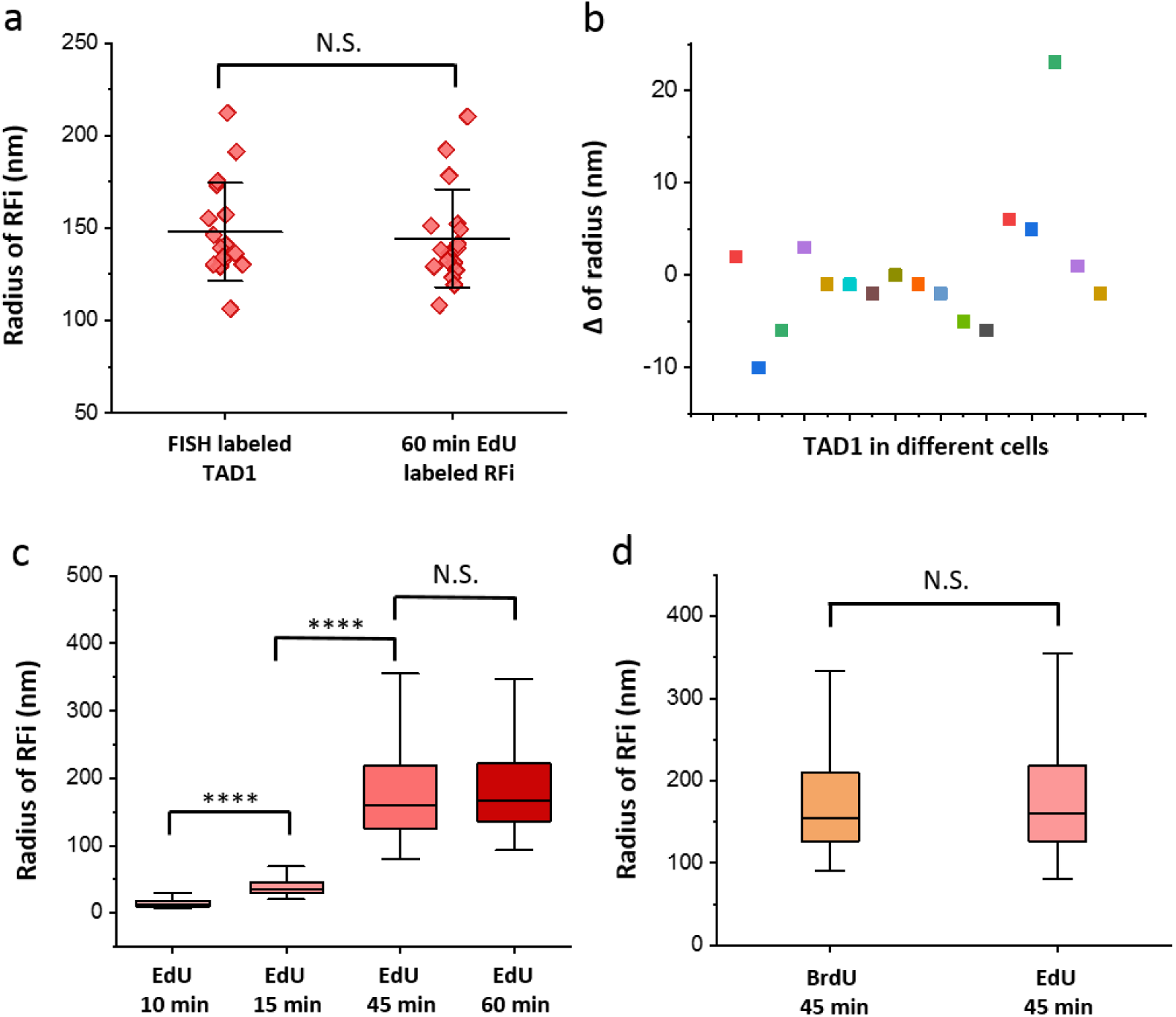
Quantitative characterization of metabolically labeled RFi. **a & b,** Comparison of FISH-labeled TAD1 and co-localized 60-min EdU labeled RFi. **a**, Radii of FISH-labeled TAD1 and 60-min EdU labeled RFi. **b**, Changes in radius between FISH-labeled TAD1 and its corresponding 60-min EdU labeled RFi in the same cell. Different colors represent different cells. **c**, Radii of RFi labeled for 10 min, 15 min, 45 min and 60 min upon release into the S phase. **d**, Radii of RFi labeled for 45 min by BrdU (yellow) or EdU (pink) at the beginning of the S phase. For lines and statistics in **a**, **c**, and **d** see the description in the legend of Figure 1 (n =10 cells).

**Appendix Figure S7.**
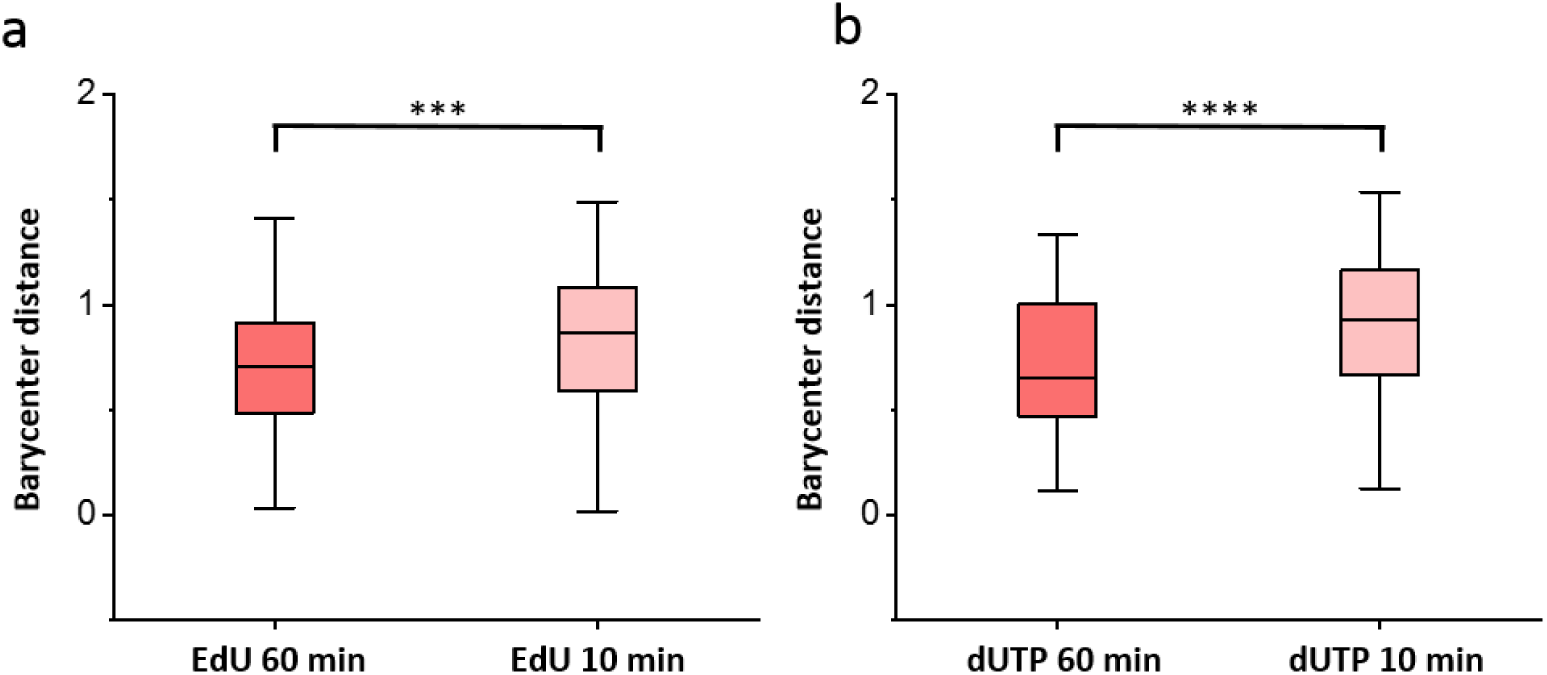
Comparison of RFi labeled by different metabolic labeling methods and for different durations. **a,** Box plot of barycenter distances between BrdU-labeled RFi and EdU-labeled RFi. BrdU was supplied for 45 min upon release into the S phase, whereas EdU was supplied for 10 or 60 min. **b,** Box plot of barycenter distances between EdU-labeled RFi and dUTP-atto550 labeled RFi. EdU was supplied for 45 min upon release into the S phase, whereas dUTP was supplied for 10 or 60 min. For lines and statistics in **a** and **b** see the description in the legend of Figure 1 (n =10 cells).

**Appendix Figure S8.**
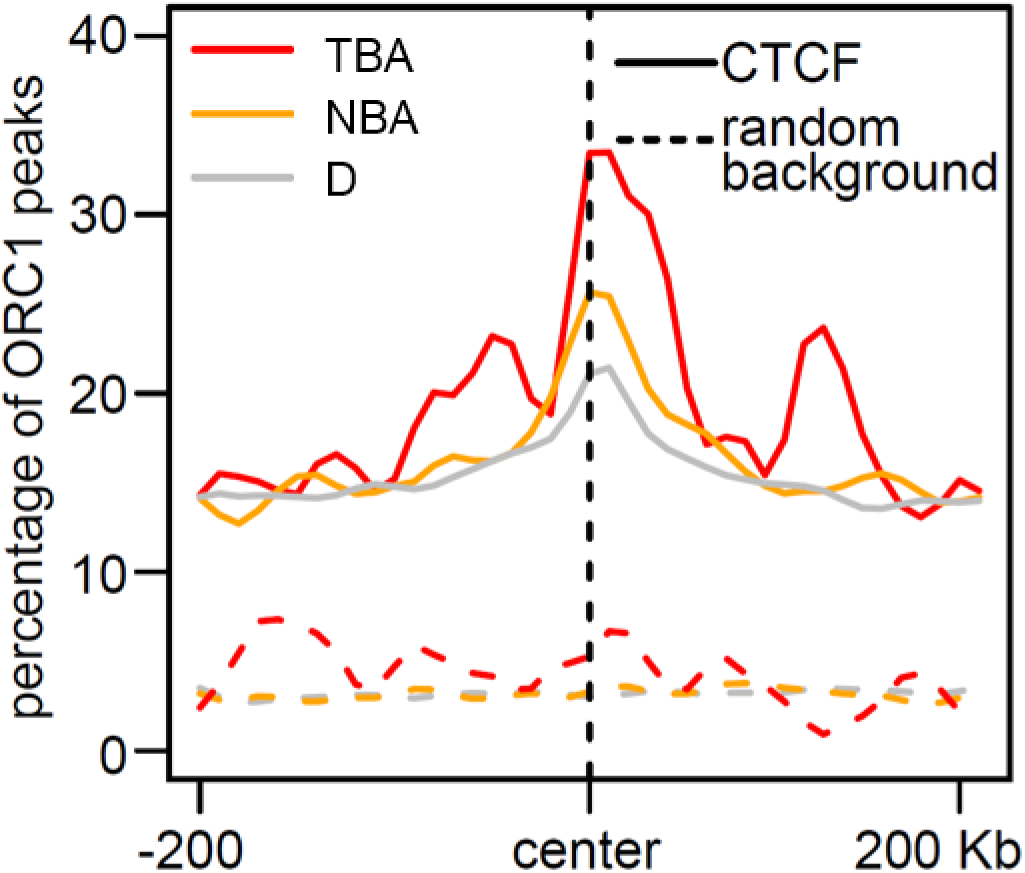
Co-localization of replication origins with CTCF-cohesin binding sites. Histograms of distance between CTCF-cohesin binding sites and replication origins are presented in solid lines; those for randomly-selected sites are presented by dotted line. TBA (TAD boundary active origins): red lines; NBA (Non-TAD boundary active origins): yellow lines; D (dormant replication origins): grey lines. Center dashed line is the sites with overlapped binding of CTCF and cohesin.

**Appendix Figure S9.**
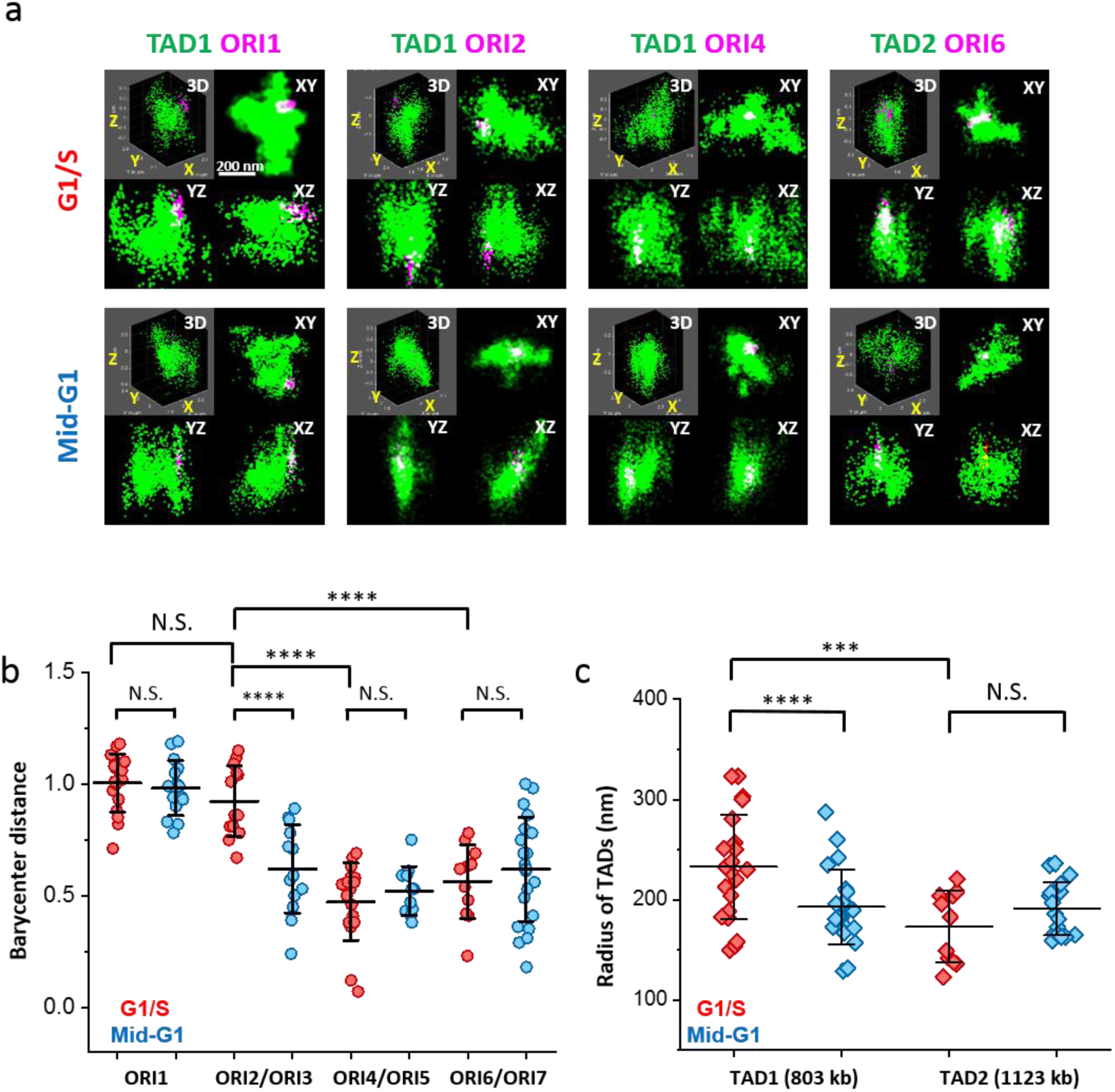
3D distribution of replication origins in TADs in the G1 and G1/S phase. The definitions and labeling procedures of TADs and origins are identical with those in Figure 2. **a**, Representative 3D STORM images of TADs (green) and their origins (purple) in the G1 and G1/S phases. Upper row: TADs and origins labeled at the G1/S transition. Lower row: TADs and origins labeled approximately 5 hours into the G1 phase. **b**, 3D Barycenter distances between all 7 origins and the 2 related TADs in a. **c,** 3D radius of gyration of TAD1 and TAD2 in the G1 and G1/S phases. For lines and statistics in **b** and **c** see the description in the legend of Figure 1 (n ≥10 cells).

**Appendix Figure S10.**
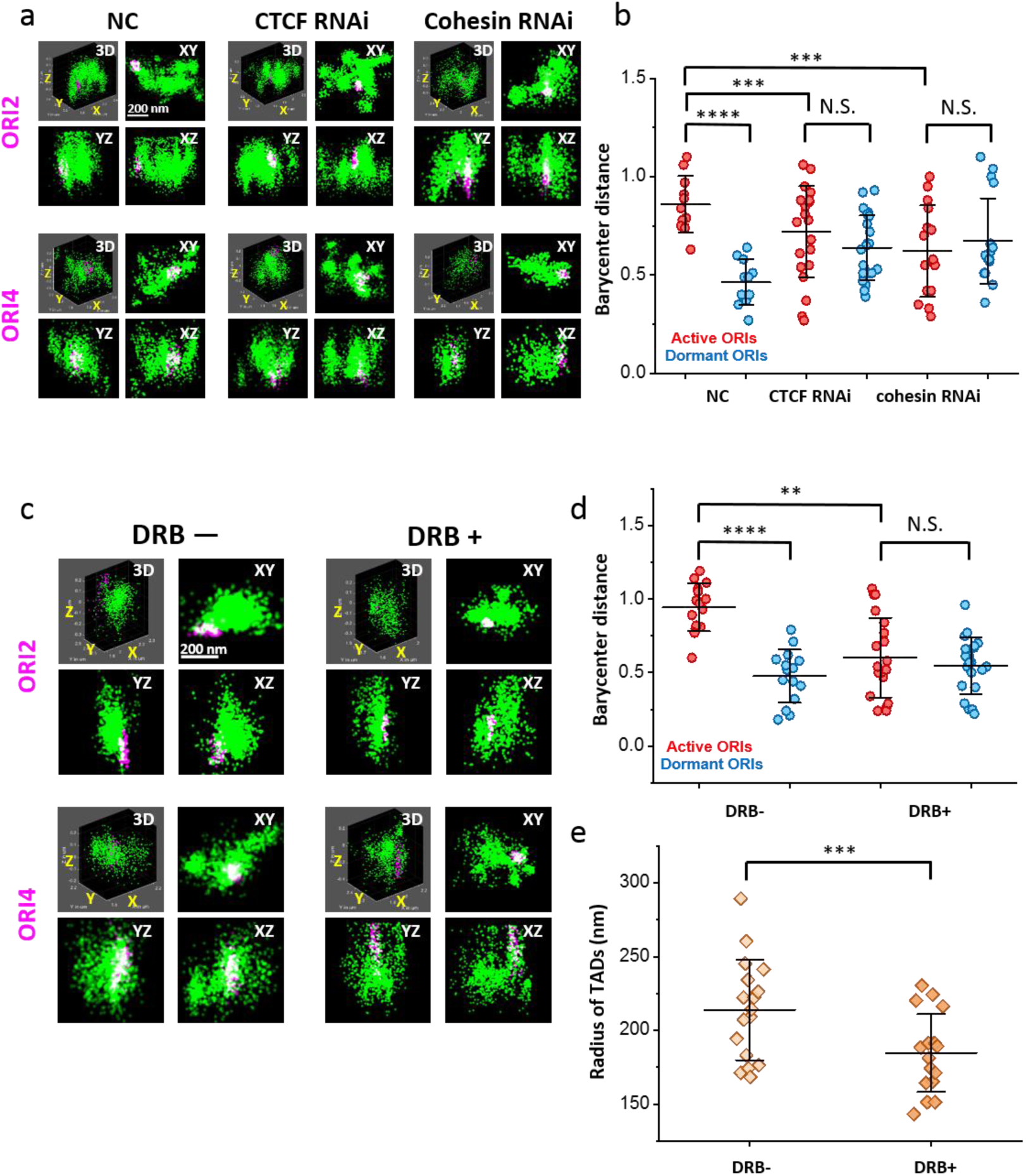
3D Distribution of replication origins in TAD1 with transcription elongation inhibition or down-regulation of CTCF or cohesin. **a,** Representative 3D STORM images of origins (purple) in TAD1 (green) after treatment of cells with the indicated siRNAs. **b**, 3D barycenter distances between active (ORI2 and ORI3) or dormant (ORI4 and ORI5) origins in TAD1 after treatment of cells with the indicated siRNAs as in **a**. **c**, Representative 3D STORM images of origins (purple) in TAD1 (green). Restricted by the space, only ORI2 and ORI4 are shown. Left: no DRB. Right: with DRB. **d,** 3D barycenter distances between active (ORI2 and ORI3) and dormant (ORI4 and ORI5) origins in TAD1 with or without DRB treatments. **e**, 3D radius of gyration of TAD1 treated with or without DRB. For lines and statistics in **b**, **d**, and **e** see the description in the legend of Figure 1 (n ≥10 cells).

**Appendix Figure S11.**
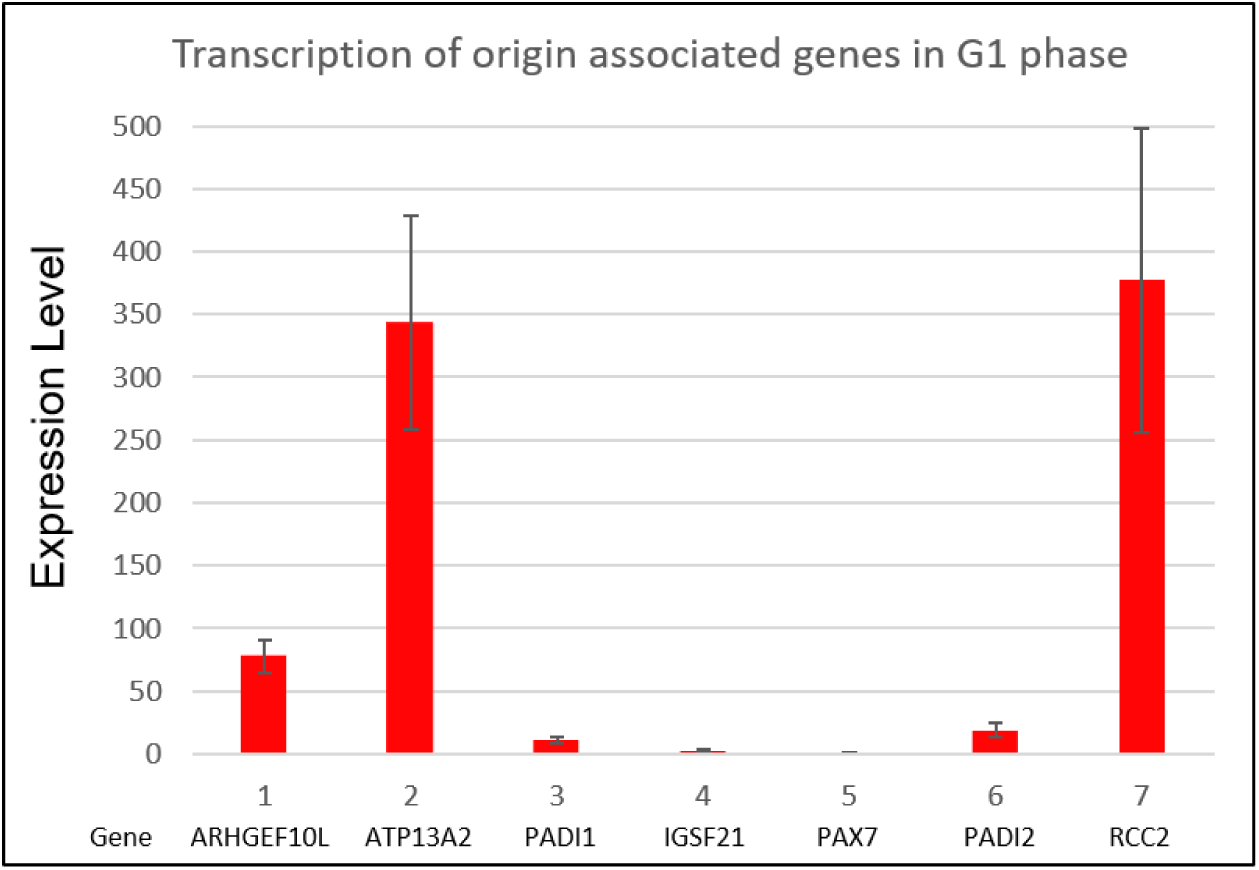
Transcription of origin-associated genes in the G1 phase. The expression data of the genes in TAD1 and TAD2 were obtained from the RNA-seq data set in the NCBI Gene Expression Omnibus (GEO; http://www.ncbi.nlm.nih.gov/geo/) under accession number GSE73565. The data clearly reveal that in TAD1, the expression level of genes associated with active replication origins (ORI1, ORI2, ORI3) are several folds of that associated with dormant replication origins (ORI4, ORI5) (3 replicates). We note that while the active origin (ORI7) the late replicating TAD2 is not exposed to the domain periphery in the early S phase, its associated gene RCC2 is actively expressed in the G1 phase. This observation supports the model that origin firing is subordinate to regulated replication timing of RDs [4, 5].

**Appendix Figure S12.**
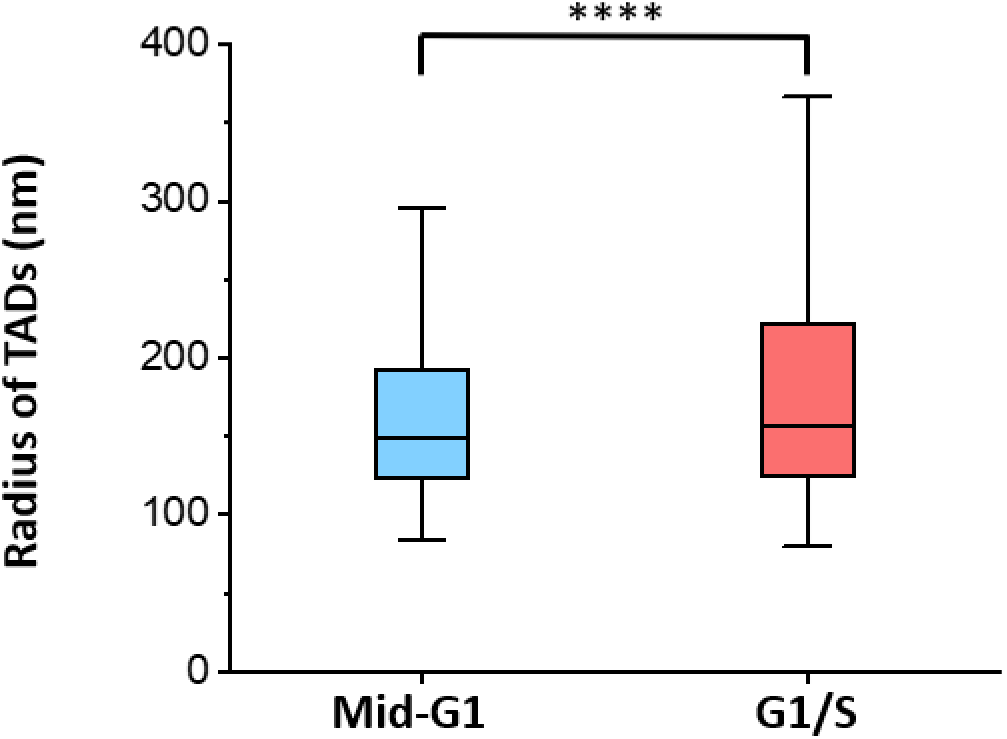
Radii of metabolically labeled TADs in the G1 and G1/S phase. TADs were labeled by EdU for 45 min upon release into the S phase. In the next cell cycle, cells were fixed in the mid-G1 or G1/S phase. For lines and statistics see the description in the legend of Figure 1 (n =10 cells).

**Appendix Figure S13.**
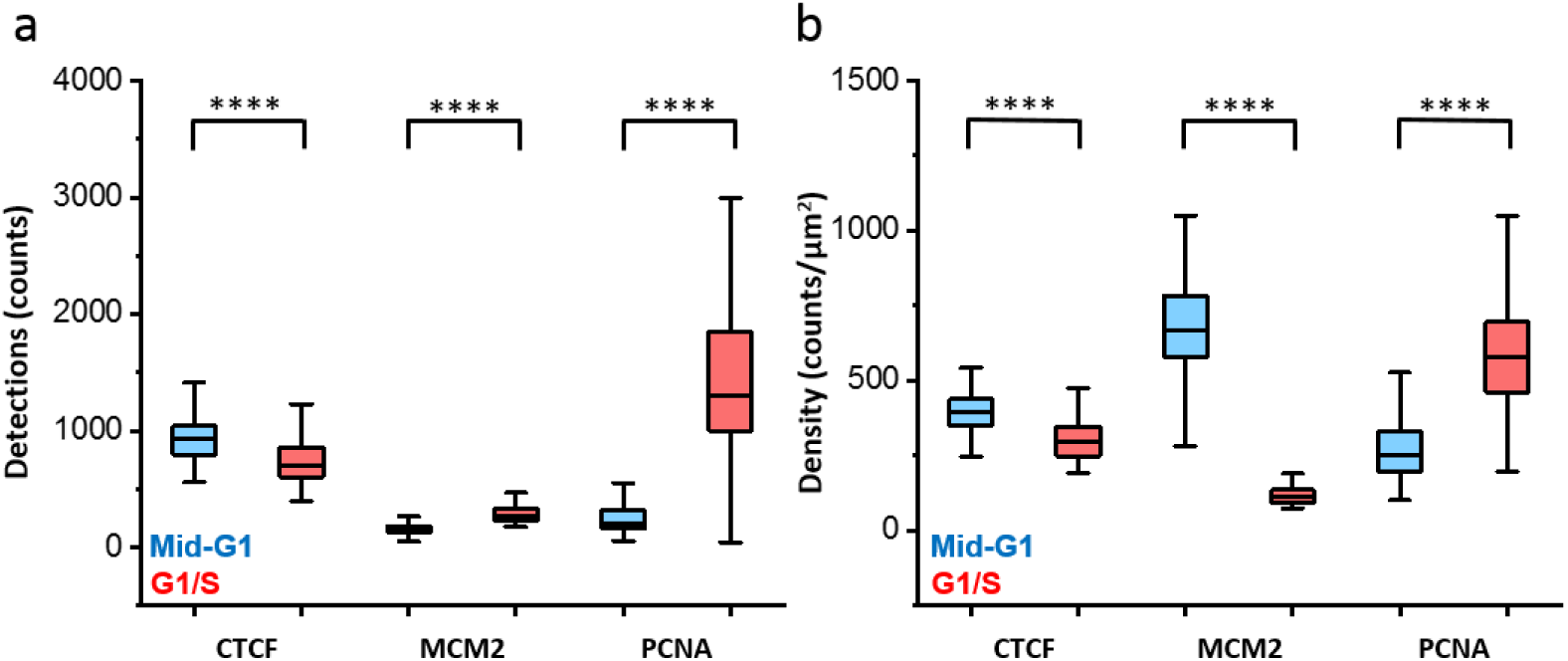
Single molecule detection counts and density of CTCT, MCM2, and PCNA in early replicating TADs in the mid-G1 and G1/S phases. Counts are shown in **a** and molecule density is shown in **b**. The reduced number of single-molecule CTCF detections and molecule density in the CTCF foci indicate that CTCF molecules dissociate from chromatin in the G1 phase. The increased number of single-molecule MCM2 detections and decreased molecule density in the MCM2 foci indicate gradual association of MCM2 with chromatin and dislocation from DNA. The increased number of single-molecule PCNA detections and molecule density in the PCNA foci indicate assembly of replication factories in the G1 phase. See details in the main text. For lines and statistics see the description in the legend of Figure 1 (n =10 cells).

**Additional File Table 1.**
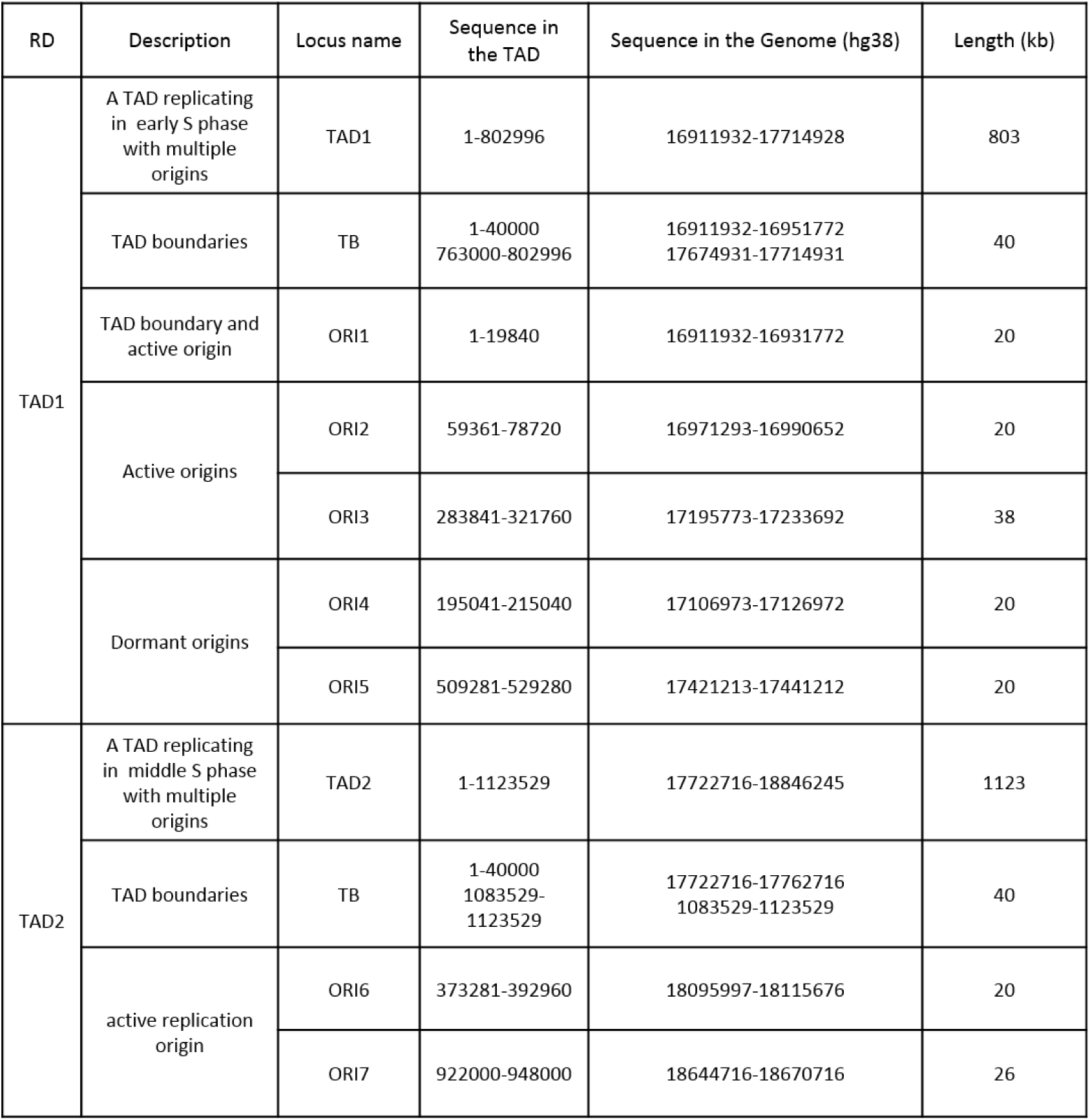
Information for TADs and origins.

