## Supplementary material for "Transcription-coupled structural dynamics of topologically associating domains regulate replication origin efficiency": supplenmental files

#### Supplementary Figures and Figure Legends

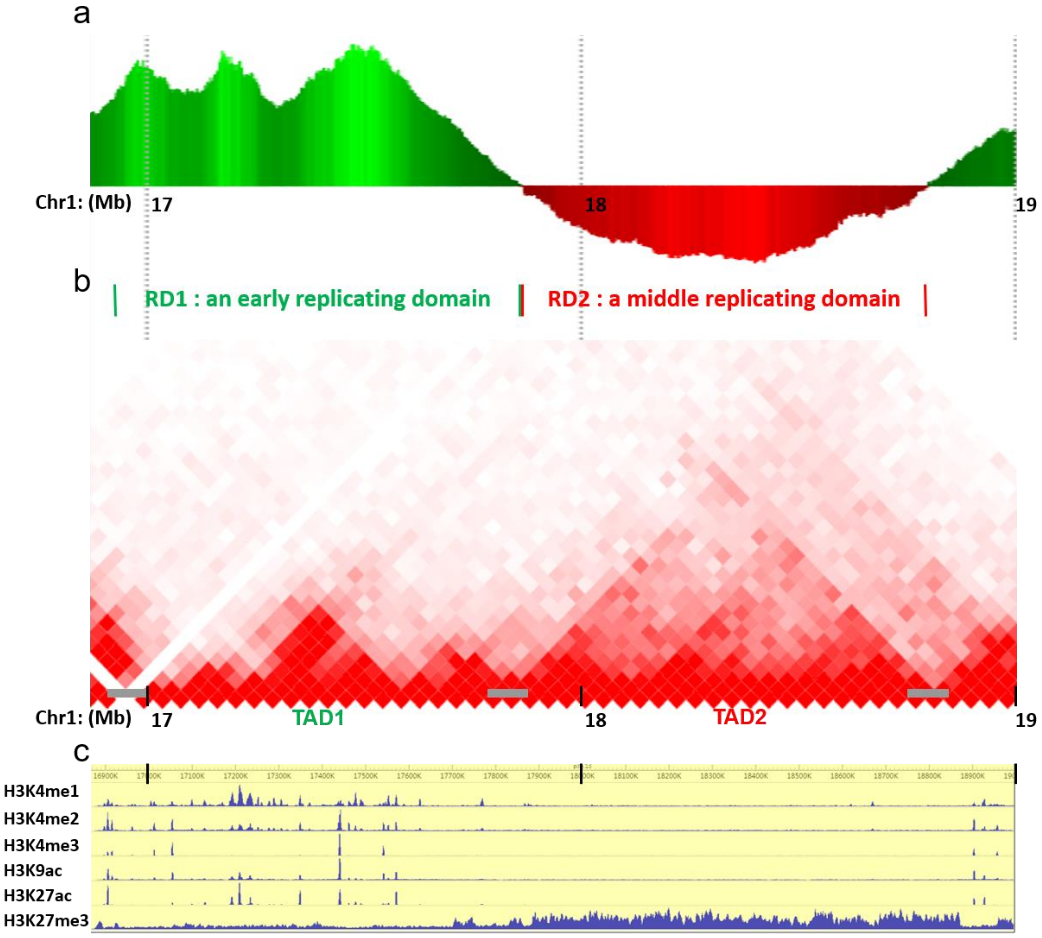


**Appendix Figure S1 Identification and histone modifications of two TADs from the replication timing profile and Hi-C interaction heatmap of HeLa cells. a-b,** Depiction of the replication timing profile (green peaks for early RDs and red peaks for middle or late RDs) and Hi-C interaction heatmap in the same genomic region of chromosome 1. The replication timing profile was obtained from the Replication Domain Genome Browser of the Gilbert lab (https://www2.replicationdomain.com/_genome_browser). The Hi-C interaction heatmap was obtained from ENCSR693GXU. Grey bars: TAD boundaries. (Methods) Two TADs were selected. TAD1: an early replicating domain (Chr1:16911932-17714928). TAD2: a middle replicating domain (Chr1:17722716-18846245). **c,** Profiles of histone modifications are from public data hubs (ENCODE data portal) of WashU Epigenome Browser.

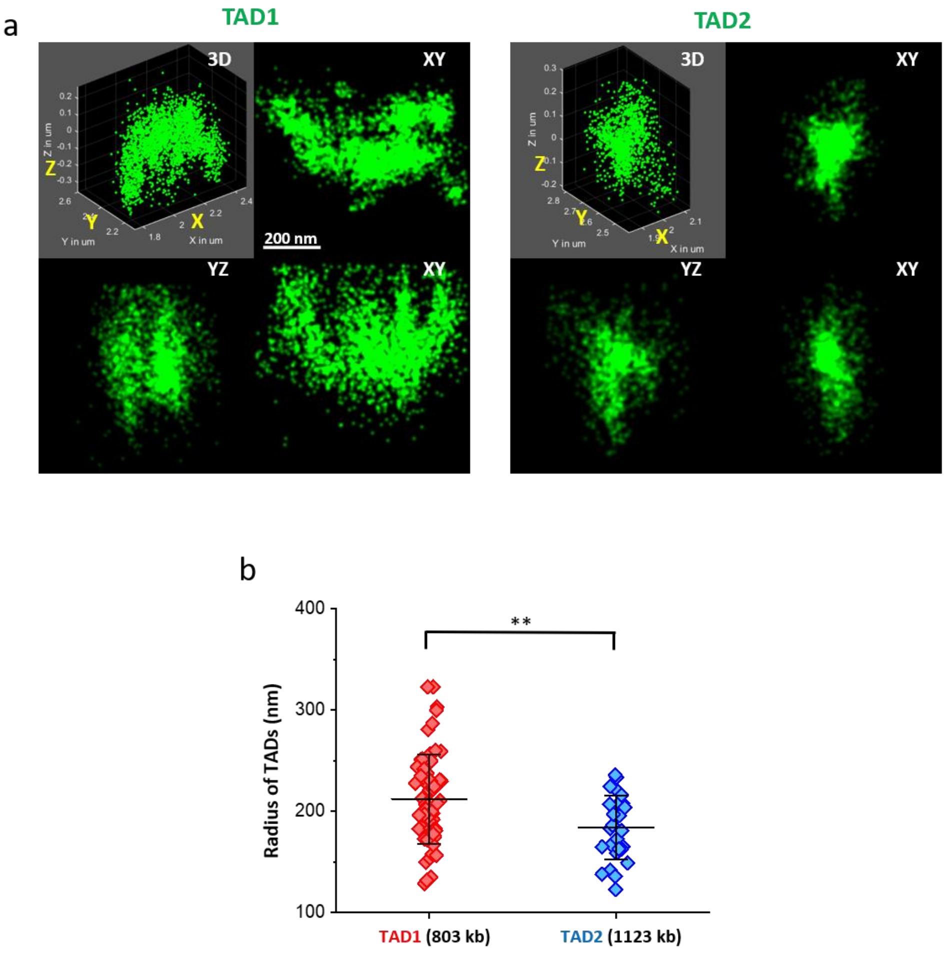


**Appendix Figure S2. 3D visualization and radii of TAD1 and TAD2.** The definitions and labeling procedures of TADs and origins are identical with those in Figure 2. **a**, Representative 3D STORM images of TADs. One 3D presentation with 3 projected images. **b,** 3D radius of gyration of TAD1 and TAD2. For lines and statistics in **b** see the description in the legend of Figure 1 (n ≥20 cells).

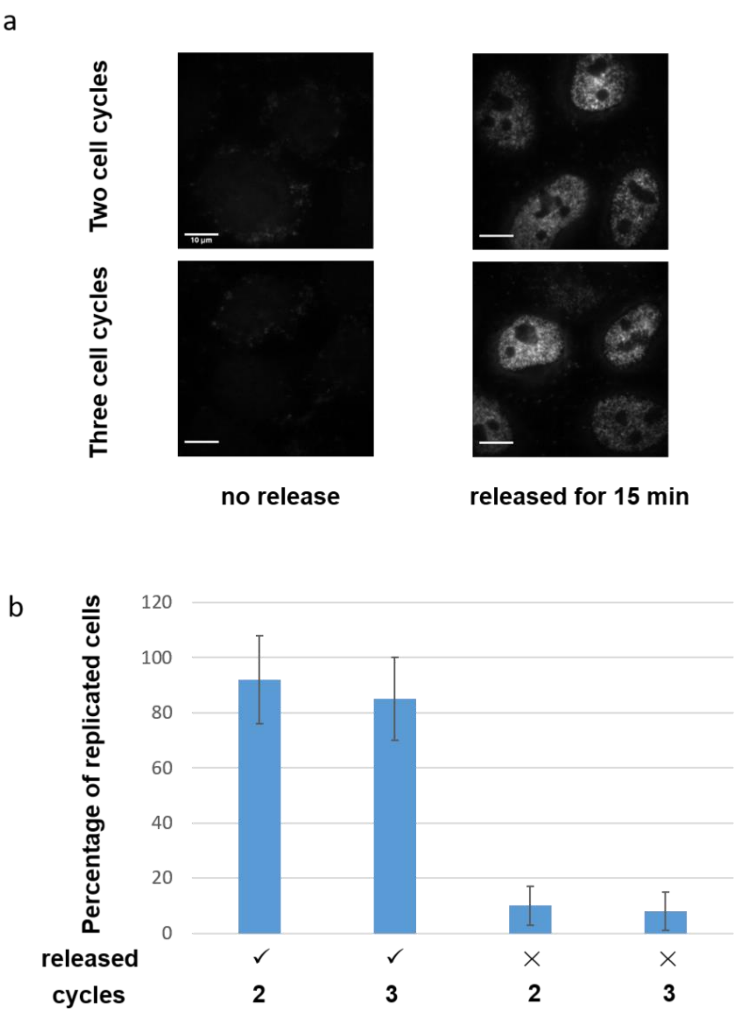


**Appendix Figure S3. Quantification of cell synchronization by EdU labeling.** To make sure the cells were successfully synchronized and the synchronization procedure minimally impacts the growth and morphology of cells, we imaged and analyzed the cells synchronized for two or three cell cycles. **a**, after two (upper) or three (lower) cycles of synchronization, cells were synchronized to the G1/S transition. EdU were added when the synchronized cells were released (right) for 15 min or were not released (left). b, percentage of the replicated cells in **a**. Replicated cells were defined by the three folds of the fluorescence of the nucleus to the background (3 replicates, 200 cells for each group). EdU labeling showed more than 80% cells entering the S phase, similar with the previous work [1].

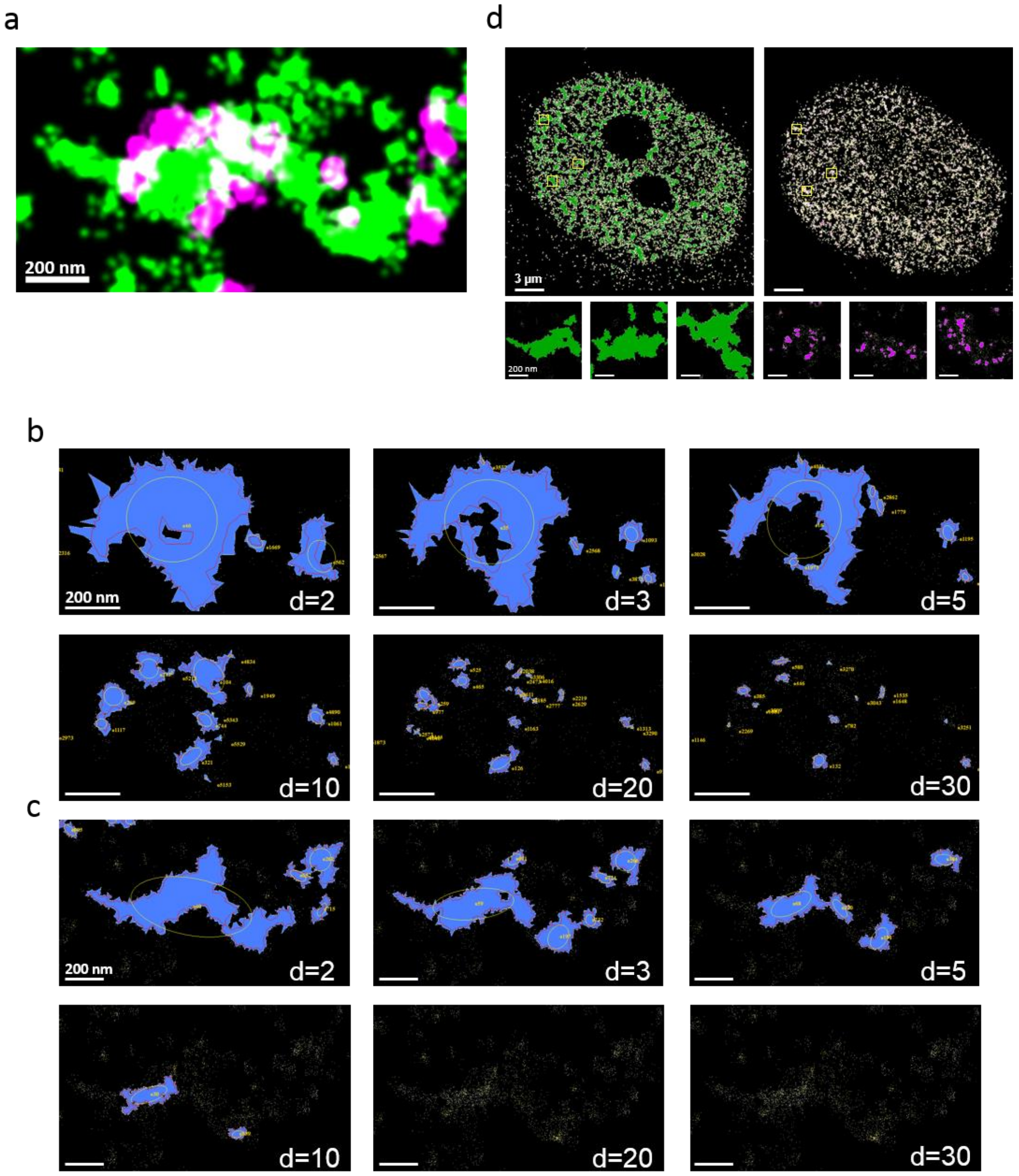


**Appendix Figure S4. Quantification of the density factor in SR-Tesseler analysis. a,** Dual-color STORM imaging of TADs (also early replicating domains, green) and origins (purple) as in Fig. 1c. **b** and **c,** Analysis of TADs and origins by SR-Tesseler with density factors from 2 to 30. **b**, Origins close to each other in a TAD cannot be separated when the density factor is set from 2 to 10. Origins are too small or even dismissed when the density factor is set to 30. When the density factor was set to 20, approximately 5,000 origins were clearly defined at the beginning of the S phase in one cell, similar with a previous report [2]. **c**, TADs close to each other cannot be separated when the density factor is set to 2. TADs are too small or even dismissed when the density factor is set from 5 to 30. When the density factor was set to 3, approximately 700 TADs were clearly identified, similar with a previous report[3]. **d**, TADs (green) and origins (purple) identified by SR-Tesseler from the STORM images using a density factor of 3 for TADs and a density factor of 20 for origins. The areas inside the yellow squares are shown at higher magnification below each nucleus.

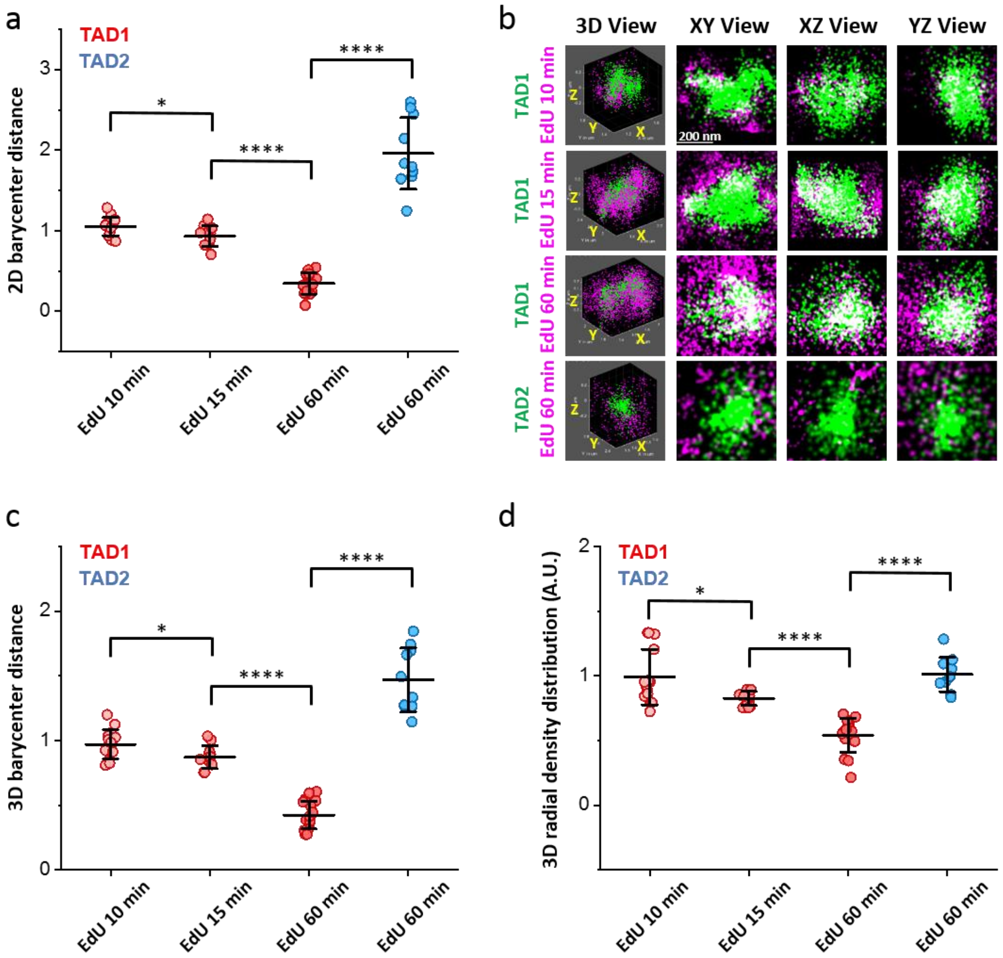


**Appendix Figure S5. Replication patterns of TADs in the S phase as determined by DBSCAN.** **a,** 2D barycenter distances between the TADs and their spatially associated RFi as determined by DBSCAN (Methods) in Figure 1**a**. **b,** Representative 3D STORM images of TAD1 and TAD2 labeled by Oligopaint probes (green) and RFi labeled metabolically for different durations (purple). Metabolic labeling of DNA replication was performed by supplying EdU to the cells upon release into the S phase for 10 min, 15 min, and 60 min (purple). **c,** 3D barycenter distances between the TADs and their spatially associated RFi as determined by DBSCAN in **b**. **d,** Radial density distribution of the RFi in TADs as determined by DBSCAN (Methods) in **b**. For lines and statistics in **a**, **c**, and **d** see the description in the legend of Figure 1 (n ≥10 cells).

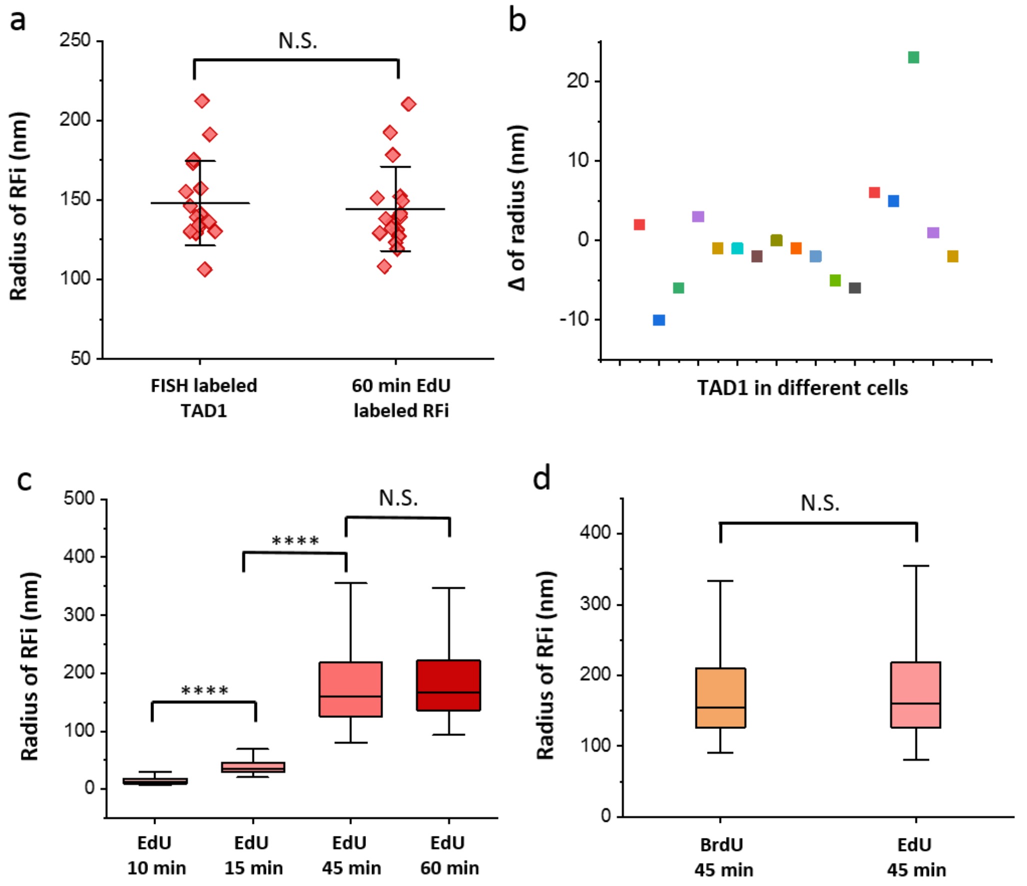


**Appendix Figure S6. Quantitative characterization of metabolically labeled RFi. a & b,** Comparison of FISH-labeled TAD1 and co-localized 60-min EdU labeled RFi. **a**, Radii of FISH-labeled TAD1 and 60-min EdU labeled RFi. **b**, Changes in radius between FISH-labeled TAD1 and its corresponding 60-min EdU labeled RFi in the same cell. Different colors represent different cells. **c**, Radii of RFi labeled for 10 min, 15 min, 45 min and 60 min upon release into the S phase. **d**, Radii of RFi labeled for 45 min by BrdU (yellow) or EdU (pink) at the beginning of the S phase. For lines and statistics in **a**, **c**, and **d** see the description in the legend of Figure 1 (n =10 cells).

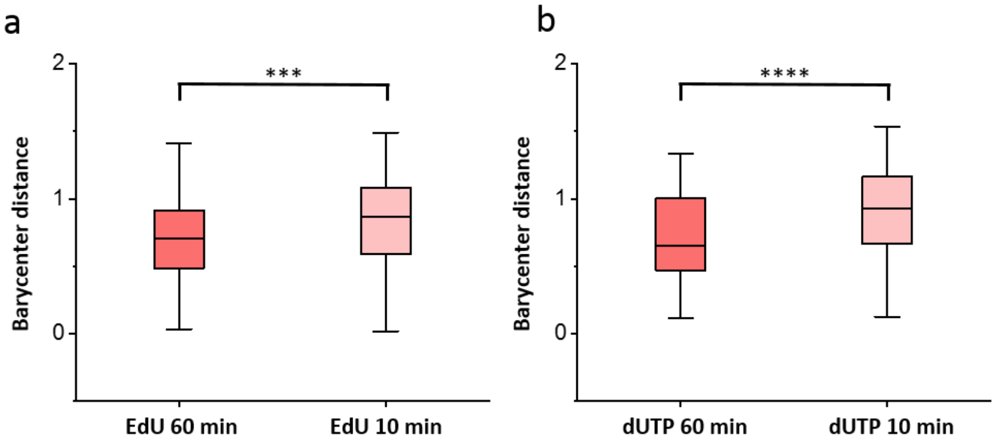


**Appendix Figure S7. Comparison of RFi labeled by different metabolic labeling methods and for different durations. a,** Box plot of barycenter distances between BrdU-labeled RFi and EdU-labeled RFi. BrdU was supplied for 45 min upon release into the S phase, whereas EdU was supplied for 10 or 60 min. **b,** Box plot of barycenter distances between EdU-labeled RFi and dUTP-atto550 labeled RFi. EdU was supplied for 45 min upon release into the S phase, whereas dUTP was supplied for 10 or 60 min. For lines and statistics in **a** and **b** see the description in the legend of Figure 1 (n =10 cells).

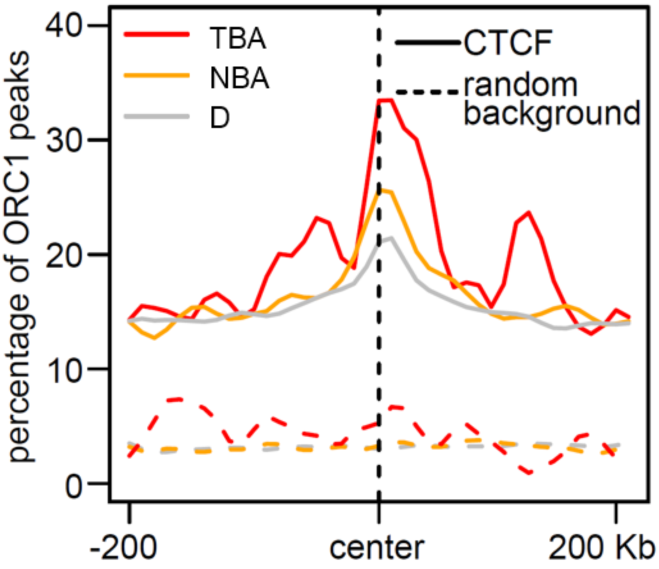


**Appendix Figure S8.** Co-localization of replication origins with CTCF-cohesin binding sites. Histograms of distance between CTCF-cohesin binding sites and replication origins are presented in solid lines; those for randomly-selected sites are presented by dotted line. TBA (TAD boundary active origins): red lines; NBA (Non-TAD boundary active origins): yellow lines; D (dormant replication origins): grey lines. Center dashed line is the sites with overlapped binding of CTCF and cohesin.

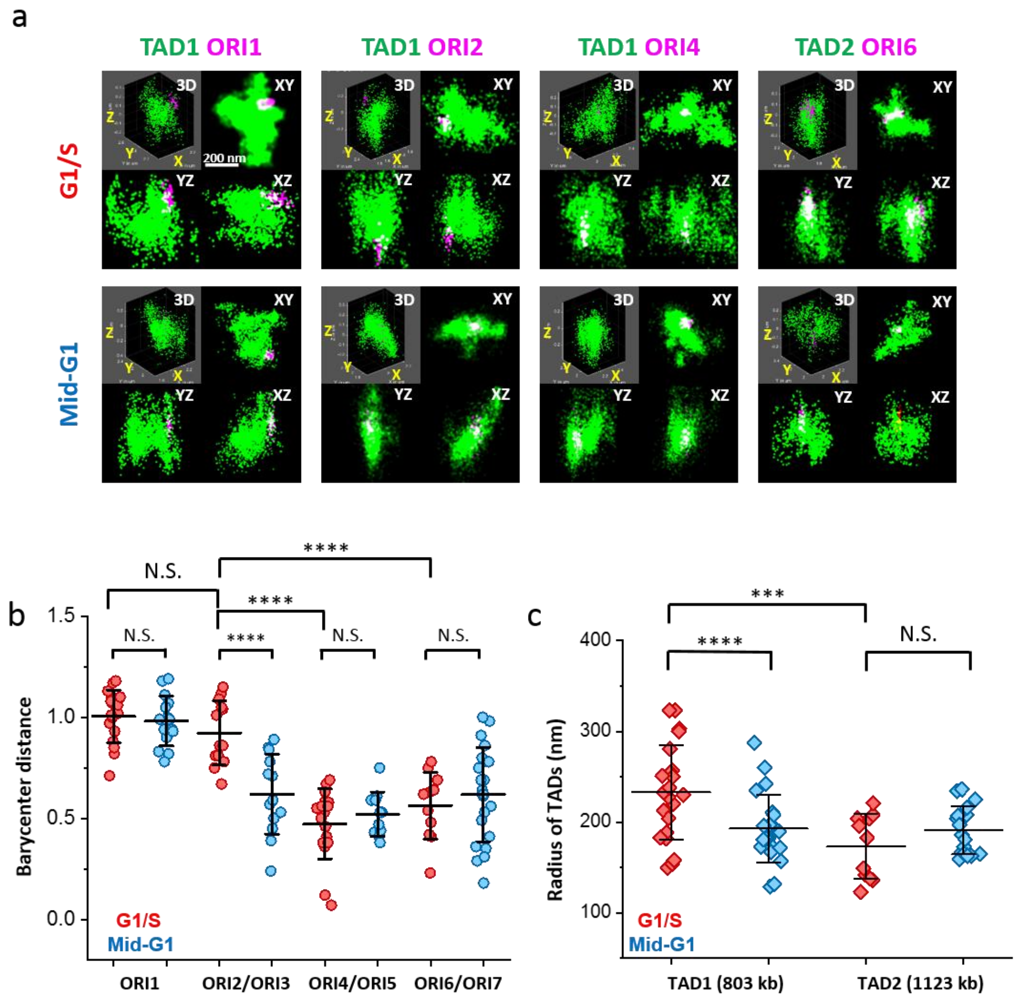


**Appendix Figure S9. 3D distribution of replication origins in TADs in the G1 and G1/S phase.** The definitions and labeling procedures of TADs and origins are identical with those in Figure 2. **a**, Representative 3D STORM images of TADs (green) and their origins (purple) in the G1 and G1/S phases. Upper row: TADs and origins labeled at the G1/S transition. Lower row: TADs and origins labeled approximately 5 hours into the G1 phase. **b**, 3D Barycenter distances between all 7 origins and the 2 related TADs in a. **c,** 3D radius of gyration of TAD1 and TAD2 in the G1 and G1/S phases. For lines and statistics in **b** and **c** see the description in the legend of Figure 1 (n ≥10 cells).

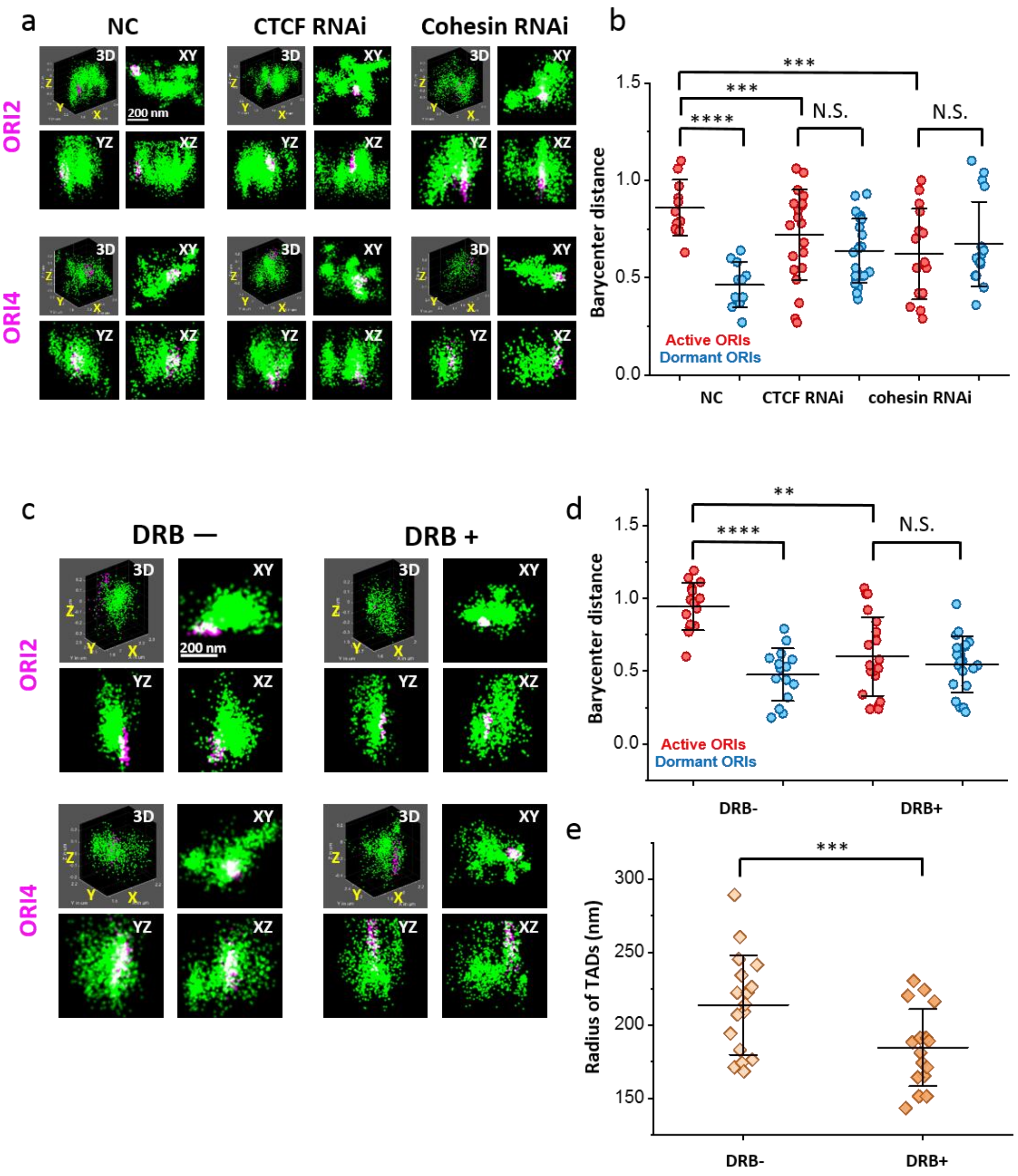


**Appendix Figure S10. 3D Distribution of replication origins in TAD1 with transcription elongation inhibition or down-regulation of CTCF or cohesin. a**, Representative 3D STORM images of origins (purple) in TAD1 (green) after treatment of cells with the indicated siRNAs. **b**, 3D barycenter distances between active (ORI2 and ORI3) or dormant (ORI4 and ORI5) origins in TAD1 after treatment of cells with the indicated siRNAs as in **a**. **c**, Representative 3D STORM images of origins (purple) in TAD1 (green). Restricted by the space, only ORI2 and ORI4 are shown. Left: no DRB. Right: with DRB. **d,** 3D barycenter distances between active (ORI2 and ORI3) and dormant (ORI4 and ORI5) origins in TAD1 with or without DRB treatments. **e**, 3D radius of gyration of TAD1 treated with or without DRB. For lines and statistics in **b**, **d**, and **e** see the description in the legend of Figure 1 (n ≥10 cells).

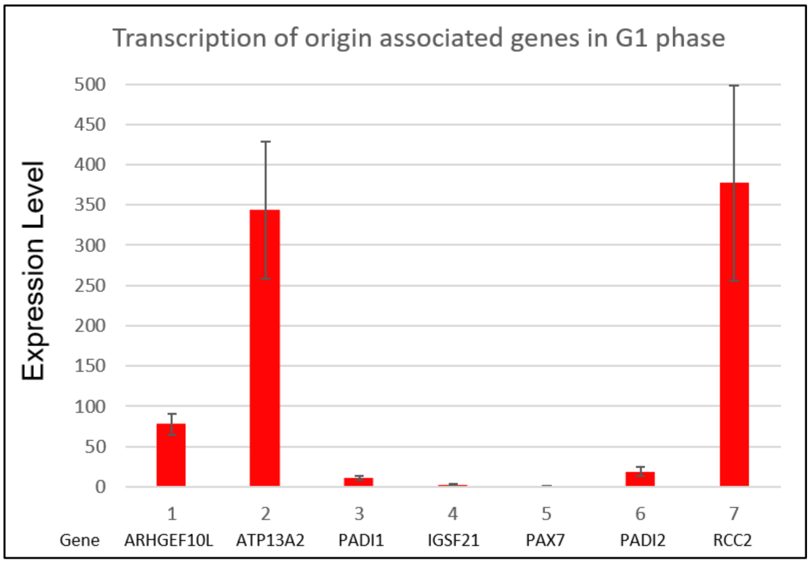


**Appendix Figure S11. Transcription of origin-associated genes in the G1 phase.** The expression data of the genes in TAD1 and TAD2 were obtained from the RNA-seq data set in the NCBI Gene Expression Omnibus (GEO; http://www.ncbi.nlm.nih.gov/geo/) under accession number GSE73565. The data clearly reveal that in TAD1, the expression level of genes associated with active replication origins (ORI1, ORI2, ORI3) are several folds of that associated with dormant replication origins (ORI4, ORI5) (3 replicates). We note that while the active origin (ORI7) the late replicating TAD2 is not exposed to the domain periphery in the early S phase, its associated gene RCC2 is actively expressed in the G1 phase. This observation supports the model that origin firing is subordinate to regulated replication timing of RDs [4, 5].

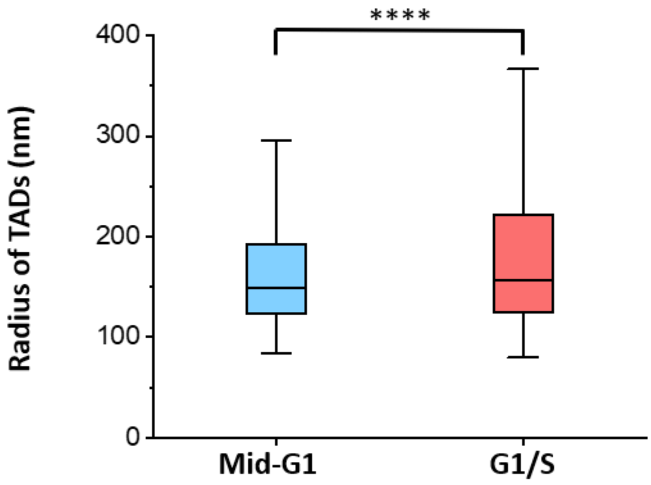


**Appendix Figure S12. Radii of metabolically labeled TADs in the G1 and G1/S phase.** TADs were labeled by EdU for 45 min upon release into the S phase. In the next cell cycle, cells were fixed in the mid-G1 or G1/S phase. For lines and statistics see the description in the legend of Figure 1 (n =10 cells).

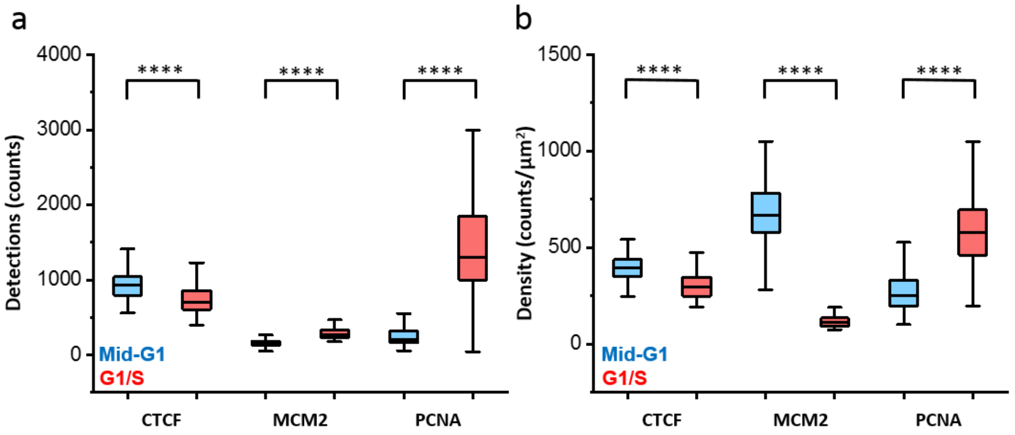


**Appendix Figure S13. Single molecule detection counts and density of CTCT, MCM2, and PCNA in early replicating TADs in the mid-G1 and G1/S phases.** Counts are shown in **a** and molecule density is shown in **b**. The reduced number of single-molecule CTCF detections and molecule density in the CTCF foci indicate that CTCF molecules dissociate from chromatin in the G1 phase. The increased number of single-molecule MCM2 detections and decreased molecule density in the MCM2 foci indicate gradual association of MCM2 with chromatin and dislocation from DNA. The increased number of single-molecule PCNA detections and molecule density in the PCNA foci indicate assembly of replication factories in the G1 phase. See details in the main text. For lines and statistics see the description in the legend of Figure 1 (n =10 cells).

Additional File Table 1 Information for TADs and origins.

| RD | Description | Locus name | Sequence in the TAD | Sequence in the Genome (hg38) | Length (kb) |
|---|---|---|---|---|---|
| TAD1 | A TAD replicating in early S phase with multiple origins | TAD1 | 1-802996 | 16911932-17714928 | 803 |
| | TAD boundaries | TB | 1-40000<br>763000-802996 | 16911932-16951772<br>17674931-17714931 | 40 |
| | TAD boundary and active origin | ORI1 | 1-19840 | 16911932-16931772 | 20 |
| | Active origins | ORI2 | 59361-78720 | 16971293-16990652 | 20 |
| | | ORI3 | 283841-321760 | 17195773-17233692 | 38 |
| | Dormant origins | ORI4 | 195041-215040 | 17106973-17126972 | 20 |
| | | ORI5 | 509281-529280 | 17421213-17441212 | 20 |
| TAD2 | A TAD replicating in middle S phase with multiple origins | TAD2 | 1-1123529 | 17722716-18846245 | 1123 |
| | TAD boundaries | TB | 1-40000<br>1083529-1123529 | 17722716-17762716<br>1083529-1123529 | 40 |
| | active replication origin | ORI6 | 373281-392960 | 18095997-18115676 | 20 |
| | | ORI7 | 922000-948000 | 18644716-18670716 | 26 |

### Methods

**Cell culture and manipulations**

The HeLa-S3 cell line (PubMed ID: 5733811) was obtained from Dr. Wei Guo, University of Pennsylvania. Hela S3 cells were grown in 10 mm glass-bottom imaging dishes (Cellvis #D35-10-1-N) with 2 mL modified medium (high-glucose DMEM, Thermo Fisher Gibco #10569-044) supplemented with 10% (v/v) fetal bovine serum (Thermo Fisher Gibco #10091-148) and 100 U/mL penicillin-streptomycin antibiotics (Thermo Fisher Gibco #15070-063) under regular cell culture conditions (37 ℃, 5% CO2, humidified atmosphere). Cells were passaged with the proportion of 1:8-1:10 by trypsin (Life Technologies #25200-056) every 3 days or when they reached 80% confluence.

Hela cells were synchronized to the G1/S boundary by two rounds of blocking as described previously [3] [6, 7]. In the first blocking period, HeLa cells at approximately 25% confluence were treated with 2 mM thymidine (sigma #T1895-5G) for 15 hours and then released to fresh medium for 10 hours. In the second blocking period, cells were arrested by 2 μg/mL aphidicolin (sigma #A0781-1MG) for 15 hours.

For FISH labeling of the TADs and metabolic labeling of replicating DNA in Fig. 1a, synchronized cells were labeled by EdU for the designated durations upon release into the S phase, followed by Click reactions and FISH labeling. For double metabolic labeling of replicating DNA in Fig. 1c, synchronized cells were labeled by BrdU for 45 min and then transferred to fresh medium. Cells were then synchronized by aphidicolin, labeled with EdU for the designated durations upon release into the next S phase, and fixed. For double FISH labeling of the TADs and origins at the G1/S transition in Fig. 2b, synchronized cells were released with fresh medium and synchronized again until they were fixed at the next G1/S phase for FISH labeling. For double FISH labeling of the TADs and origins at the mid-G1 phase in Fig. 2b, cells were synchronized in mitosis as previously described [8]. In brief, after release from the aphidicolin block, cells were transferred to fresh medium and nocodazole was added to a final concentration of 0.1 μg/mL. After incubation for 10 hours, mitotic cells were collected by mechanical shaking and transferred to fresh medium. After 7 hours, cells were fixed at 5 hours in the G1 phase, *i.e.* mid-G1 phase, for FISH labeling. For Fig. 3a, after down-regulation of proteins, cells were synchronized and fixed at the G1/S transition for double FISH labeling. For Fig. 3c, synchronized cells were released, after which DRB was added at a final concentration of 100 μM from 5 hours in the G1 phase to the S phase in the third cell cycle. Cells were fixed at the G1/S transition for double FISH labeling. For metabolic labeling of early replicating TADs and immunolabeling of proteins the beginning of the S phaseat the beginning of the S phase in Fig. 4, synchronized cells were labeled by EdU for 45 min and then transferred to fresh medium. Cells were synchronized again until they were fixed at the next G1/S phase for protein immunolabeling. For metabolic labeling of TADs and immunolabeling of proteins at the mid-G1 phase in Fig. 4, synchronized cells were labeled by EdU for 45 min and then synchronized in mitosis as previously described [8]. In brief, after release from the aphidicolin block, cells were transferred to fresh medium and nocodazole was added to a final concentration of 0.1 μg/mL. After incubation for 10 hours, mitotic cells were collected by mechanical shaking and transferred to fresh medium. After 7 hours, cells were fixed at 5 hours in the G1 phase, *i.e.* mid-G1 phase, for protein immunolabeling.

For the Extended Data Figures, cells were manipulated using the same methods used for the experiments shown in the related figures. The procedures used for FISH and immunolabeling are described in detail below.

**Metabolic replication labeling**

At specific times after releasing the cells from the G1/S boundary, replication foci were labeled by directly adding thymidine analogs EdU (Click-iT® Plus Alexa Fluor® 647 Picolyl Azide Toolkit, Thermo Fisher Invitrogen #C10643) or BrdU (Abcam #ab142567) for a designated time period. The final concentration of EdU and BrdU was 10 μM. dUTP-atto-550 (Jena Bioscience #NU-803-550-S) was delivered into the nucleus using the FuGENE 6 transfection reagent (Promega #E2691) according to the manufacturer's instructions. BrdU was further labeled by immunostaining as described in detail below. Alexa Flour 647 was conjugated by click reaction according to the manufacturer's instructions. In brief, after fixation and washing, samples were treated with 1% Triton in 5% BSA for 30 min. The samples were washed three times, after which fresh reaction cocktail was added for 30 min. The reaction cocktail was made by sequentially mixing 1× Click-iT reaction buffer, copper protectant, Alexa Fluor picolyl azide and reaction additive buffer. Finally, the samples were washed three times.

**Sequencing data**

The replication timing profile remapped to hg38 was obtained from the Replication Domain Genome Browser of the Gilbert lab (https://www2.replicationdomain.com/ genome_browser). Hi-C data were obtained from ENCSR693GXU and further analyzed using previously described methods [9]. In brief, Hi-C data was analyzed using HiCUP (v0.5.8) using default parameters, then duplicates were removed using samtools (v1.0.0). Analyzing HiC from the Homer suite (v4.8) was used for the coverage (-window 1000000) and DLR (distal-to-local ratio) calculation (-dlrDistance 3000000). For BrdU-seq, in brief, cells were synchronized to the beginning of the S phase and released, after which BrdU was supplied for 10 min at 0 min, 1 hour, 3 hours and 6 hours into the S phase. Sample preparations and data analyses for the

ChIP-seq experiments were performed following standard protocols [10]. ChIP-seq data sets of CTCF and cohesions (SMC1 and SMC3) was initially analyzed as described for histone ChIP-seq data analysis. Summits of the CTCF peaks overlapped with cohesion peaks were remained for the distribution analysis of ORC1 peaks. Sequencing data was remapped to hg38.

#### Replication domain labeling by fluorescence *in situ* hybridization

##### *Probe design*

To label target genomic regions, corresponding oligonucleotide probes were designed using Oligominer (https://github.com/brianbeliveau/Oligominer) following online instructions and methods described in a previous work [11]. All templates for primary hybridization probes synthesis were 102 nt in length and contained the following components: i) one 32 nt central sequence targeting the genomic region of the target, ii) one 30 nt flanking sequence to hybridize with the secondary probes, iii) two 20 nt flanking primer binding sequences to amplify the probes through polymerase chain reaction (PCR). For probe synthesis, a 20 nt T7 promoter sequence was added to the 5′ end of each forward primer for *in vitro* transcription. The primers used for probe synthesis and the secondary probe sequences are listed below.

**Additional File Table 2 Information for probes and primers.**

| Primer | Sequence |
|---|---|
| TAD1-primary primer F with T7 | GCCGTACGGATAATACGACTCACTATAGGG CCCGCGTTAACCATACACCG |
| TAD1-primary primer R | CATCGAAGCGTGTGGCTACC |
| TAD1-reverse transcription primer | CATCGAAGCGTGTGGCTACC(5’TAMRA modified) |
| Origin 1 | CGCAACGCTTGGGACGGTTCCAATCGGATC (5′ AF647 modified) |
| Origin 2 | CGAATGCTCTGGCCTCGAACGAACGATAGC (5′ AF647 modified) |
| Origin 3 | AAGTCGTACGCCGATGCGCAGCAATTCACT (5′ AF647 modified) |
| Origin 4 | CAAGTATGCAGCGCGATTGACCGTCTCGTT (5′ AF647 modified) |
| Origin 5 | ACGAATCCACCGTCCAGCGCGTCAAACAGA (5′ AF647 modified) |
| TAD2-primary primer F with T7 | GCCGTACGGATAATACGACTCACTATAGGG GTGGTAAAGCTCCGCGGCTT |
| TAD2-primary primer R | TCGTTCCGCATTGACCAATC |
| TAD2-reverse transcription primer | TCGTTCCGCATTGACCAATC (5′ TAMRA modified) |
| Origin 6 | CGCGCGGATCCGCTTGTCGGGAACGGATAC (5′ AF647 modified) |
| Origin 7 | GCCCGTATTCCCGCTTGCGAGTAGGGCAAT (5′ AF647 modified) |

***Probe synthesis***

The probes were amplified from a template oligopool synthesized by Hongxun Biotech (Suzhou, China). All primers were synthesized by Invitrogen. The probe synthesis was performed by a five-step enzymatic amplification procedure as follows:

Step 1: Through 26 cycles PCR with the above primers, 2 μL of the template oligopool was amplified in a 50 μL reaction mix (Phanta Max Super-Fidelity DNA Polymerase, #P505-d2). The final PCR product was column-purified (Zymo DNA Clean and Concentrator, DCC-5).

Step 2: To obtain the target products at a high concentration, the purified DNA products were collected and used as the template to repeat Step 1. Next, the purified PCR products were diluted to a concentration of 100 ng/μL.

Step 3: The diluted DNA was converted into RNA *via in vitro* transcription (HiScribe™ T7 High Yield RNA Synthesis Kit NEB, #E2040S). Each 33 μL reaction contained 17 μL of template DNA obtained as described above, 2.5 μL (10 mM) of each NTP, 3 μL of 10× reaction buffer, 1 μL of RNase inhibitor (Promega RNasin® Plus RNase Inhibitor #N2615) and 2 μL T7 polymerase. The reaction was incubated at 37 °C for 5 to 6 hours.

Step 4: RNA products from the *in vitro* transcription reaction were converted back into single stranded DNA *via* reverse transcription (MAN0012047 TS Maxima H Minus Reverse Transcriptase, Thermo Fisher #EP0751). Each 70 μL reaction contained 33 μL of the RNA products obtained in Step 3, 10 μL dNTPs, 14 μL RT buffer, 1 μL RT enzyme, 1 μL RNase inhibitor (Promega RNasin® Plus RNase Inhibitor #N2615), 1 μL ddH2O and 10 μL 40 μM TAMRA-labeled reverse transcription primer. The reaction was incubated at 50 °C for 1.5 hours.

Step 5: The template RNA obtained in Step 4 was removed by incubating it with 50 μL of 0.25 M EDTA and 0.5 M NaOH at 95 °C for 10 min. Next, the products were put on ice and immediately purified by column purification (Zymo Research, #D4006). The resulting single-strand DNA probes were eluted twice in 20 μL of ultra-pure water. Probes purified in this manner can be stored temporarily at -20 °C or at -80 °C for several months. The quality of the probes was determined by measuring the sample concentration (200–300 ng/μL) and the fluorescence absorbance of tagged dyes.

***In situ hybridization***

Step 1: Sample preparation

Cultured cells were prepared for fluorescence *in situ* hybridization (FISH) as described in previous works [11, 12]. Briefly, cells were fixed on dishes with 4% paraformaldehyde, washed three times in PBS, and incubated in 1 mg/mL $NaBH_4$ solution for 7 minutes to quench background signals. If FISH is combined with other labeling methods, such as immunostaining or click-reaction, FISH should be performed after other labeling steps, and the quenching step must be skipped. Cells were then permeabilized with PBS containing 1% Triton X-100 (PBST) for 10 min and washed twice in the same buffer. To remove all RNA, cells were treated with 100 μg/mL RNaseA in 1× PBST for 45 min at 37 ℃, after which they were washed three times in PBST. The samples were incubated with 0.1M HCl in 1× PBST for 10 min, followed by two washes in 2× SSCT. To unfold chromatin, 50% formamide in 2× SSCT was added to the samples for at least 4 hours or overnight.

Step 2: Fluorescence *in situ* hybridization

The cells were incubated with 50% formamide in 2× SSCT at 78 °C for 10 min and then put on ice immediately. Next, the samples were dehydrated in 70%, 85% and 100% icy ethanol successively for 1 minute each. Meanwhile, 5 μL of the synthesized primary probes and 1 μL of the synthesized secondary probes (100 μM) were mixed with 35 μL of 100% formamide at 37 °C in a 1500 rpm vortex for 15 min. At the same time, 20% Dextron/4× SSC was shaken at 37 °C in a 1500 rpm vortex for 15 min. Next, 41 μL of formamide with probes and 35 μL of 20% Dextron/4× SSC added by 1 μL of triton were mixed and shaken for 30 min at 37 °C in a 1500 rpm vortex. The fluorescent probes were kept in darkness. The 77 μL hybridization cocktail was incubated at 86 °C for 3 min and immediately put on ice before the hybridization. The processed sample was incubated with the hybridization cocktail at 86 °C for 3 min, followed by hybridization in a wet box at 37 °C for 16–20 hours.

Step 3: Post-hybridization washes

The samples were briefly washed three times with 2× SSCT. Next, the samples were washed three times with 2× SSCT at 60 °C for 10 min. The signal and background noise of the samples were checked by microscopic imaging. If the background noise was high, the samples were washed several times with 2× SSCT and 50% formamide at 37 °C for 1 min each time to obtain an appropriate signal-to-noise ratio. Finally, the sample was stored in 2× SSCT prior to STORM imaging. All reagents mentioned in this section of the methods were purchased from Sigma.

**Immunostaining of BrdU, PCNA, MCM and CTCF**

Cells were fixed by 4% PFA for 15 min and then blocked and permeabilized in 5% bovine serum albumin (BSA) containing 1% Triton X-100 (Sigma-Aldrich) for 30 min, after which the cells were washed three times in PBS containing 1% (v/v) Triton. All primary antibodies (CTCF,

Abcam #ab128873; PCNA, CST #13110T; MCM, CST #3619T; BrdU, Abcam #ab152095) were diluted (1:200) in 5% BSA containing 1% Triton. Next, cells were incubated with primary antibodies for 1 hour in darkness, followed by three washes in PBS (5 min, 10 min, and 5 min). Cells were incubated with 1:50 dye-conjugated secondary antibody for 1 hour followed by four washes in PBS (5 s, 5 min, 10 min, and 5 min). After the cells were post-fixed in 4% PFA for 10 min, they were washed three times with PBS (15 seconds each) and stored at 4 °C. All of the steps described above were performed at room temperature.

**RNA interference**

siRNA oligos were transfected into cells with Lipofectamine 2000 (Invitrogen #11668-027) at a final concentration of 100 nM according to the manufacturer's instructions. As previously reported [13], two rounds of transfection were separated by 24 h. The first round of interference was conducted when the cells reached approximately 30% confluence. Cells were released to fresh medium after the transfection, which lasted for 5 hours. After approximately 20 hours, RNA interference was conducted again when the cells reached approximately 50% confluence. After the cells were released to fresh medium for approximately 4 hours, thymidine was added and two rounds of synchronization were performed as described above. Control transfections were performed with scrambled control siRNA. siRNAs targeted to CTCF and Rad21 were designed as follows:

CTCF: GGAGCCUGCCGUAGAAAUUTT (sense)
AAUUUCUACGGCAGGUCCTC (anti-sense)

Rad21-1: GGUGAAAAUGGCAUUACGGTT (sense)
CCGUAAUGCCAUUUUCACCTT (anti-sense)

Rad21-3: GACAUGUUAGUAAGCACUACUACUU (sense)
AAGUAGUAGUGCUUACUAACAUGUC (anti-sense)

Rad21-1 and Rad21-3 were combined in equal proportions.

Control: CGUACGCGGAAUCUUCGATT (sense)
UCGAAGUAUUCCGCGUACGTT (anti-sense)

The efficiency of siRNA knockdown in individual cells was checked by immunostaining.

**STORM imaging**

STORM Imaging was performed on a custom-built inverted microscope (IX83, Olympus) using wide-field excitation mode as previously reported [14]. The microscope utilized a 100×, 1.40 NA oil-immersion objective lens (UPlanSApo 100×, 1.40 NA, Olympus). A multiband dichroic filter (Chroma) was used to reflect the lasers while ensuring transparency to the fluorescence of the sample. Extra emission filters (Chroma) were added to separate the fluorescence in each channel, respectively. Images were recorded with an EMCCD (Ixon+ EMCCD, Andor) at 256 × 256 pixels (160 nm per pixel).

An extra NIR laser was used to achieve perfect focusing. This weak laser beam was separated by a 50:50 prism, after which two mirrors reflected the resulting beams back to the prism for merging into the same direction. After these beams were collimated to the objective from opposite edges, they were reflected back and focused onto a NIR camera, resulting in two separated points. The objective was mounted on a nanopositioning system (Nano F100S, Mad City Labs), with which the distance change from two infrared points leaded by the sample drift that could be feed-back controlled.

For 3D imaging, a cylindrical lens was inserted between the microscope and the camera to achieve optical aberration. Before imaging, a calibration curve was generated by imaging individual dye molecules while scanning the sample in the *z* direction.

Silicon dioxide beads (3 μm diameter) were incubated with the sample for 0.5–2 hours (1:500 ratio), followed by three washes with PBS to remove unstable beads. Next, the PBS was replaced with imaging buffer (10% (w/v) glucose, 20 mM NaCl, 100 mM Tris/HCl, 600 μg/mL glucose oxidase, 60 μg/mL catalase, 1% (v/v) beta-mercaptoethanol). The dish was placed on the objective stage after the microscope was configured. With the appropriate focal plane, the targeted cells were chosen based on three features: qualified infrared focal locking signal, clear images in the 561 nm and 647 nm channels, and at least one stable bead along with the cell in a bright field.

Fluorophores were activated by a 405-nm laser (OBIS, Coherent) and excited by a 561-nm laser and 647-nm laser (MBP laser), respectively. At first, cells were chosen and conventional images were collected with a 647-nm or 561-nm solid-state laser. Next, the immobile silicon dioxide beads were imaged in a bright field for the drift correction process. The probes were turned rapidly to the dark state by a 561-nm or 647-nm solid-state laser, after which they were imaged with 561-nm or 647-nm laser excitation (1000 mW, >3 kW/cm$^2$) and weak 405-nm laser activation (<2 W/cm$^2$, increased from zero manually, maintaining a roughly constant density of activated molecules). STORM imaging movies were obtained with 100 Hz imaging speed and consisted of 60,000 frames in total for the 647-nm and 561-nm channels, respectively. Interlaced bright field images of the 561-nm and 647-nm channels taken as described above were utilized for drift correction between these two channels.

After imaging, STORM imaging buffer was removed by three washes in PBS, after which the samples were stored in PBS at 4 ℃.

**STORM image analysis**

For single molecule localization, fluorophores were fitted and localized by Insight3 (gift from Prof. Bo Huang, UCSF) using the Gaussian fit method. Comparison of the *x* and *y* widths of a single detection with a reference calibration curve as mentioned above was used to determine the *z*-position. Drift correction and data rendering were performed using custom code (MATLAB 2016a). Bright field images of $SiO_2$ beads were used for cross-correlation to obtain the drift-correction curve [15]. 2D and 3D STORM images were further clustered by SR-Tesseler and DBSCAN, respectively.

For the SR-Tesseler analysis, software was downloaded from the authors' website http://www.iins.u-bordeaux.fr/team-sibarita-SR-Tesseler. A Voronoi diagram was created, after which clusters were identified by the proper density factors. Different density factors were evaluated for identification of TADs and small foci such as protein clusters or replication origins (Appendix Figure S2). The following information was directly exported: detection, density, major axis, diameter, barycenter of x and y. We defined the barycenter distance to describe the spatial distribution of replication foci, replication origins or proteins in TADs. Taking FISH-labeled replication origins and TADs as an example, the barycenter distance is the spatial barycenter distance between the origins and the TAD divided by the radius of the TAD. To ensure that the origins belonged to the TAD, an origin was analyzed only if the spatial barycenter distance between the origin and its nearest TAD was shorter than half of the length of the major axis of the TAD.

To assess our experimental results, we calculated the expected mean barycenter distance for randomly distributed foci in a polar coordinate representation:

$$\bar{r} = \frac{\int_0^{2\pi} d\varphi \int_0^{\pi} d\theta \int_0^{R} dr\, r^3 \cos^2\theta}{\int_0^{2\pi} d\varphi \int_0^{\pi} d\theta \int_0^{R} dr\, r^2 \cos\theta} = \frac{3\pi}{16}R$$

Where $\bar{r}$ denotes the average spatial distance between small foci within the TAD and $R$ represents the sampling radius, which, in our case, equals half the length of the major axis of the TAD. With normalization, the mean barycenter distance between randomly distributed foci within the TAD is 0.71.

For DBSCAN analysis, DBSCAN clustering was performed using custom Python code. In brief, for each localization, we first calculated the total localization number *N* within a threshold distance *r* from it. We set these two thresholds based on the total localization number from each image. Localizations were labeled as "core point" if their value of *N* were above the threshold. The "core points" were then clustered when their distance was under the reference distance $r_{ref}$. Then, for each cluster, "border points" were defined as other points inside the radius $r_{ref}$ from the "core points". Finally, all points from each cluster were defined and saved for further analysis. The barycenter of each cluster was calculated by averaging the coordinates of all points. The radius of gyration ($R_g$) was calculated as follows:

$$R_g^2 = \frac{1}{N}\sum_{i=1}^{N}(r_i - \bar{r})^2$$

In this formula, $\bar{r}$ is the centroid of all *N* localizations and $r_i$ is the vector of an individual localization.

The radical distribution function (RDF) is a physical parameter describing the radical statistical probability distribution, which is defined as follows:

$$\int_0^{\infty} dr * F_{RDF}(r) = 1$$

where *r* is the radical distance away from a given center and $F_{RDF}(r)$ represents the RDF. To consider the distribution tendency along the radical direction, the volume of the sample space should be normalized. We define the radical density distribution function (RDDF) for this purpose:

$$\int_0^{\infty} r^2 dr \int_{-\pi/2}^{\pi/2} \cos\theta \, d\theta \int_0^{2\pi} d\varphi * F_{RDDF}(r,\theta,\varphi) = 1$$

Under the above definition, $F_{RDDF}(r,\theta,\varphi)$ denotes the RDDF in polar coordinates, which represents the distribution of the replication initiation tendency along the radius from a TAD. We made the assumption that there is no cell polarity factor that influences the distribution of replication initiations, such that we obtained:

$$F_{RDDF}(r) = \frac{F_{RDF}(r)}{4\pi r^2}$$

For simple comparison, we calculated the mean-value from each TAD and normalized it by its radius of gyration ($R_g$):

$$RDD = \frac{mean(F_{RDDF}(r))}{R_g}$$

where the radical density distribution (*RDD*) is the physical parameter used for comparison in the figures.
